# Structural basis for polyglutamate chain initiation and elongation by TTLL family enzymes

**DOI:** 10.1101/2020.06.07.139295

**Authors:** Kishore K. Mahalingan, E. Keith Keenen, Madeleine Strickland, Yan Li, Yanjie Liu, Haydn L. Ball, Martin E. Tanner, Nico Tjandra, Antonina Roll-Mecak

## Abstract

Glutamylation, introduced by TTLL enzymes, is the most abundant modification of brain tubulin. Essential effector proteins read the tubulin glutamylation pattern, and its misregulation causes neurodegeneration. TTLL glutamylases posttranslationally add glutamates to internal glutamates in tubulin C-terminal tails (branch initiation, through an isopeptide bond), and additional glutamates can extend these (elongation). TTLLs are thought to specialize for initiation or elongation, but the mechanistic basis for regioselectivity is unknown. We present cocrystal structures of murine TTLL6 bound to tetrahedral intermediate analogs that delineate key active-site residues that make this an elongase. We show that TTLL4 is exclusively an initiase, and through combined structural and phylogenetic analyses, engineer TTLL6 into a branch-initiating enzyme. TTLL glycylases add glycines posttranslationally to internal glutamates, and we find that the same active-site residues discriminate between initiase and elongase glycylases. These active-site specializations of TTLL glutamylases and glycylases ultimately yield the chemical complexity of cellular microtubules.

## Introduction

The tubulin tyrosine ligase (TTL) and tubulin tyrosine ligase-like (TTLL) family of enzymes are responsible for the most abundant and evolutionarily widespread posttranslational modifications on microtubules. These involve the non-ribosomal, ATP-dependent addition of three different amino acids. Tubulin tyrosine ligase adds a tyrosine to the C-terminal glutamate in α-tubulin, while TTLLs are known to add either glutamates or glycines to internal glutamate residues in the intrinsically disordered tubulin C-terminal tails (reviewed in ^1^). Of these, glutamylation is the most evolutionarily widespread tubulin modification, found in organisms ranging from ciliates to humans; glutamylated microtubules are a hallmark of differentiated cells ^1^. The first glutamate in the glutamylation reaction is added through the formation of an isopeptide bond between the γ-carboxyl group of a tubulin-encoded glutamate residue and the amino group of the incoming glutamate. This is the branch initiation step (Box 1). Subsequent glutamates are then added from this initial branch-point glutamate to form a polyglutamate chain. This is the elongation step (Box 1). Formally, the glutamates in the elongated polyglutamate chain could be linked either through a standard peptide bond on α-carboxyl groups (α-linked elongation) or an iso-peptide bond using the γ-carboxyl group of the accepting glutamate (γ-linked elongation; **Box 1**). Both α-linked ^2^ and γ-linked glutamate chains ^3^ were reported in two separate studies using tubulin isolated from cells.

**Box 1.**
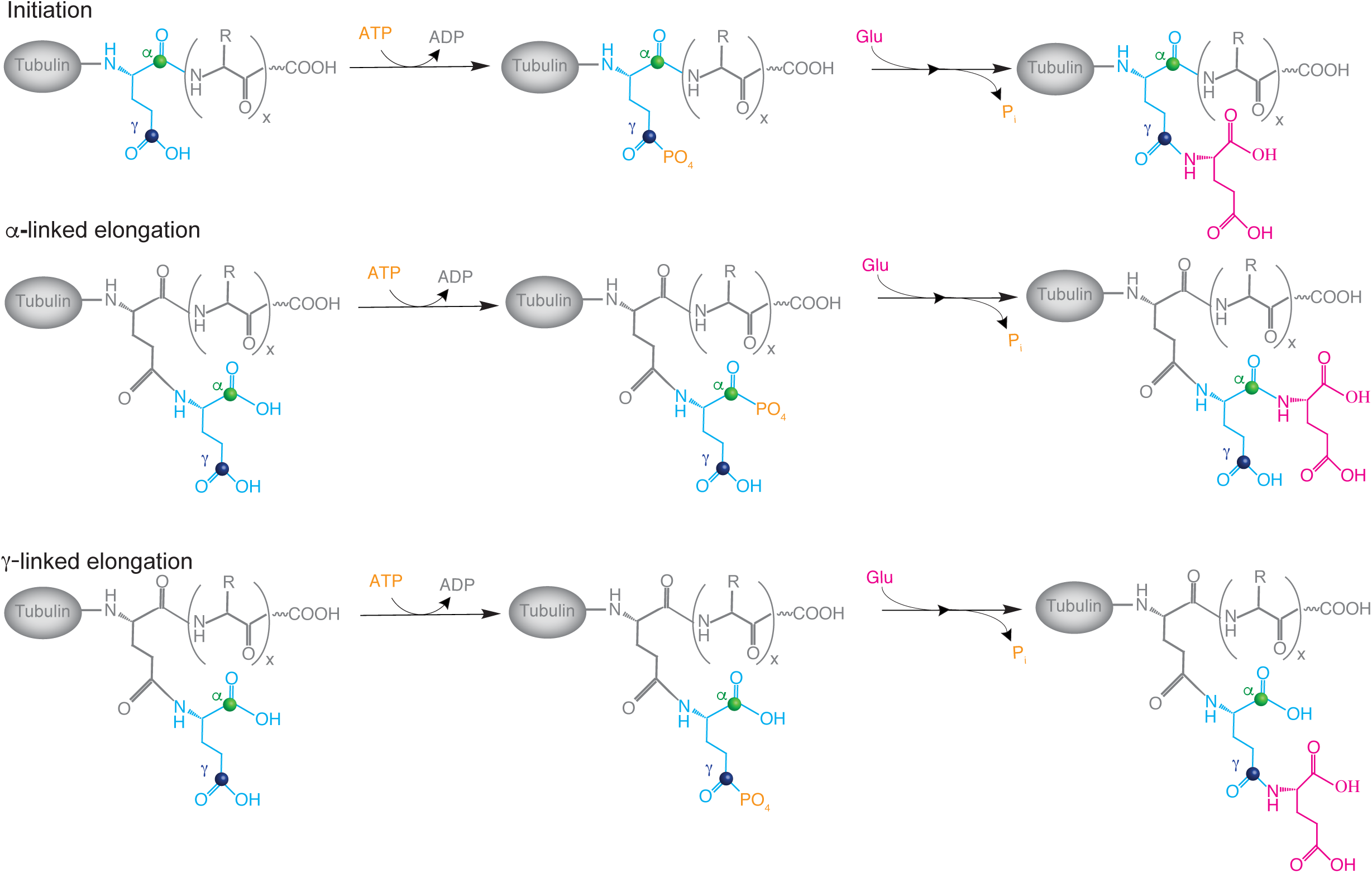
Glutamylation has two general phases: initiation and elongation. Initiation consists of the addition of a branch point glutamate to the γ-carboxyl group of an intenal glutamate in a polypeptide chain (scheme 1). Elongation consists of the addition of multiplr glutamate to a branch point glutamate. This can occur either on the α-carboxyl (scheme 2) or γ-carboxyl (scheme 3) group of the branch point glutamate.

Humans have nine TTLL family glutamylase genes ^4^. Tetrahymena, which builds sixteen different types of microtubule arrays during its lifecycle, encodes 51 TTLLs in its genome ^5^. Cellular overexpression ^4^ and biophysical studies ^6,7^ indicate that glutamylases have a preference for modifying either α-or β-tubulin and that some prefer to either initiate or elongate glutamate chains ^4^. Biochemical evidence for segregated initiation or elongation activities for purified enzymes is however lacking. Mechanistic interpretation of these overexpression studies is complicated by the apparent functional redundancy within this family as well as the variable and complex modification status of the microtubule substrate *in vivo*. Furthermore, the nature of the linkage of the glutamate chains elongated by the various TTLL glutamylases is also not known.

Microtubule glutamylation patterns are stereotyped, and both spatially and temporally regulated ^8^. The majority of tubulin in the adult nervous system is functionalized with 3-6 glutamates, with as many as 11 glutamates on α-tubulin ^9^. Microtubules in axons contain longer glutamate chains, while those in the soma and dendrites are enriched in mono-glutamylation ^10,11^. Microtubules in the axonemes of cilia are the most heavily modified, with as many as 21 posttranslationally added glutamates ^12,13^. The length of the glutamate chain on tubulin tunes the binding and activities of molecular motors ^14^ and microtubule associated proteins in a graded manner (^15,16^). Increasing evidence now supports the idea that these tubulin posttranslational modifications constitute a ‘tubulin code’ that modulates microtubules in *cis* by directly affecting microtubule dynamics and mechanical properties, and in *trans* by regulating the multitude of interactions tubulin and microtubules make with cellular effectors, and thus help microtubule networks achieve their large functional diversification in cells. Glutamylation regulates cargo transport in the axon ^17,18^. It regulates the activity of inner-arm dynein in flagella ^19,20^. It also controls the ciliary targeting of signaling complexes that are part of the hedgehog pathway by regulating anterograde intraflagellar transport ^21^. Not surprisingly, misregulation of tubulin glutamylation levels leads to neurodegenerative disorders ^17,22,23^ and ciliopathies in humans ^22,24^.

More recent studies show that glutamylation by TTLL family enzymes functions as a more general regulator of protein-protein interactions. For example, glutamylation of cyclic GMP-AMP synthase (cGAS) by TTLL6 inhibits its DNA binding, and represses the innate immune response to viral infections ^25^. Glutamylation of the retinitis pigmentosa GTPase regulator by TTLL5 is important for the targeting of opsins in photoreceptors ^26^ and its absence leads to retinal degeneration in humans ^27,28^. Glutamylation of histone chaperones by TTLL4 ^29^ is important for early embryonic development in mice ^30^, immune response to viral infections ^31^, and chromatin remodeling during pancreatic carcinogenesis in humans ^30^. Thus, understanding the catalytic activities and substrate preferences of TTLL-family members is essential to understand the broader biological functions of protein glutamylation.

Using recombinant enzymes and well-defined tubulin substrates coupled with tandem mass spectrometry (MS-MS) we show that the glutamylase TTLL4 is specialized for initiating glutamate branches on the β-tubulin tail, while TTLL6 is specialized for elongating glutamate chains on the α-tubulin tail. Using nuclear magnetic resonance (NMR) we demonstrate that the TTLL6-catalyzed glutamate chains are α-linked. High-resolution crystal structures of TTLL6 in complex with three small molecule inhibitors that are tetrahedral intermediate analogs for the initiation, and α- and γ-linked elongation reactions, respectively, reveal the structural basis for this regioselectivity. Furthermore, we identify key residues that specialize TTLL family enzymes for either initiation or α-linked elongation of glutamate chains and engineer a mutant that preferentially initiates glutamate branches as opposed to elongating them. Crystal structures of this mutant complexed with the three tetrahedral intermediate analogs reveal the structural basis for this switch in activity. Moreover, the key residue for this regioselectivity partitions between initiases and elongases for TTLL glycylases, also. Thus, our structural and functional work establishes the molecular basis for regioselectivity for the entire TTLL family.

## Results

### TTLL6 adds long glutamate chains to the α-tubulin tail

Using recombinant, purified mouse TTLL6, and synthetic tubulin tail peptides, we find that TTLL6 has low activity with either α- or β-tubulin tails, with two-fold higher activity with the α-tail (**Fig. 1a**). Interestingly, the activity of the enzyme with a detyrosinated α-tail peptide (α1B-Y), bearing a C-terminal glutamate, is eight-fold higher compared to the peptide with a C-terminal tyrosine (α1B). Given this preference of the enzyme to modify a peptide containing a glutamate with a free α-carboxylic acid, we then synthesized α-tubulin tails with a monoglutamate branch at either position 443 or 445 (**Fig. 1a**). These are reported sites of polyglutamylation on tubulin isolated from neuronal tissue ^2,32,35^. Strikingly, we see a 15- and 12-fold increase in TTLL6 activity when either E443 or E445 on the α-tail harbor a branch monoglutamate. Furthermore, peptide substrates bearing a di-glutamate branch at either of these positions further increase TTLL6 activity (**Fig. 1a**), indicating a preference for the formation of long polyglutamate chains. Mapping of the modification sites for these branched peptides using LC-MS-MS shows that chain elongations occur specifically from the preexisting branch sites and not from other glutamates in the α-tubulin tail sequence (**Fig. 1b** and **Supplementary Fig. 1**). A mono-glutamate branch also makes the β-tail peptide a better TTLL6 substrate, but the effect is not as pronounced as for the α-tubulin peptide (**Fig. 1a**), indicating that the enzyme active site has a preference for the α-tail primary sequence in addition to the branch point glutamate.

**Figure 1.**
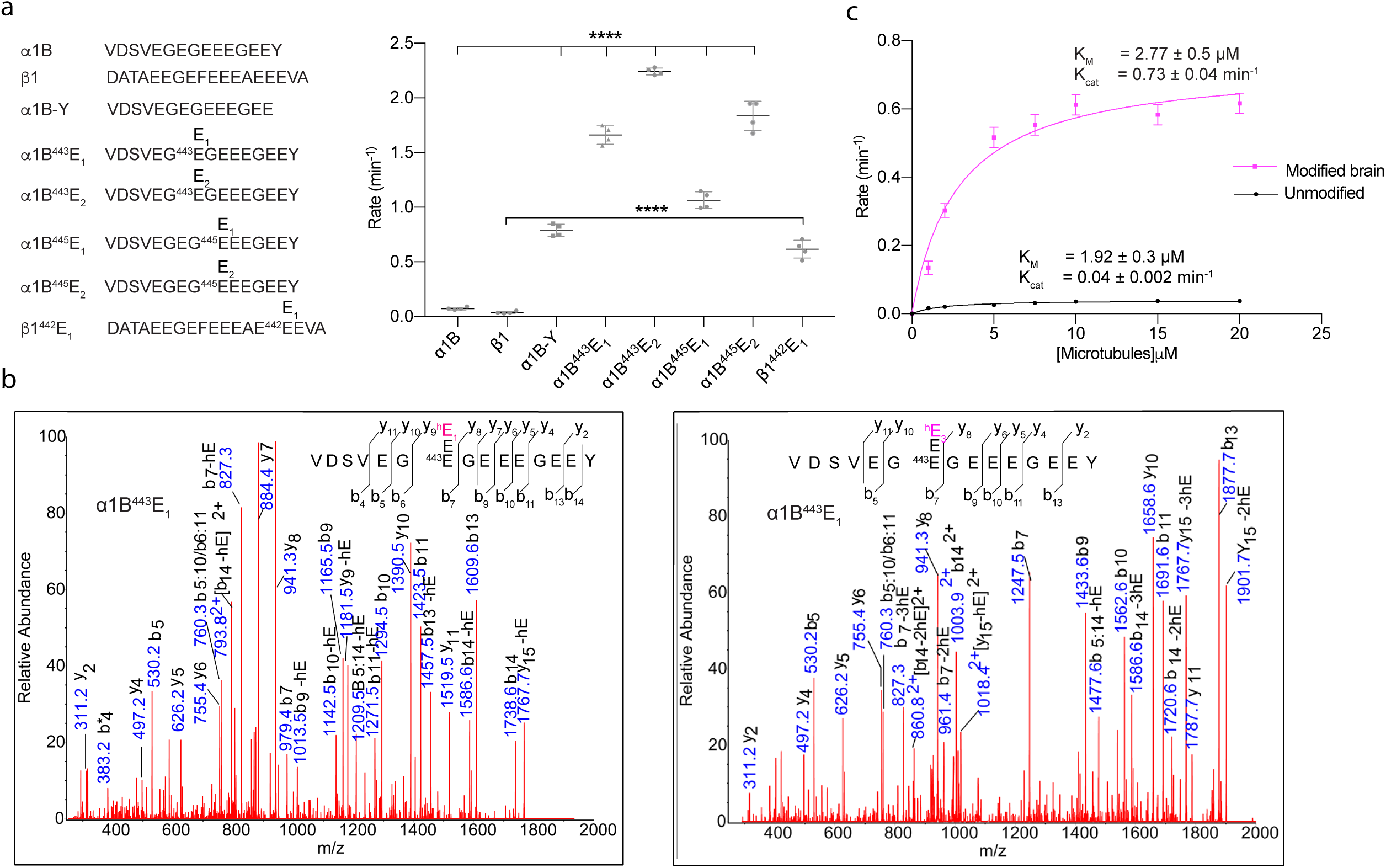
TTLL6 preferentially elongates branched glutamates in α-tubulin C-terminal tails. (**a**) Left: Sequences of C-terminal tail peptides used in this study. Branch glutamates are indicated by En (where n is the number of glutamates in the branch) over the glutamate that carries the branch point. Right: Glutamylation of the α- and β-tubulin tail peptides shown on the left by TTLL6 (n=4). Error bars indicate s.e.m. (**b**) MS-MS sequencing of the α1B^443^E_1_ peptide glutamylated by TTLL6. Species with one (left) and three enzymatically added heavy glutamates (right) added to the mono-glutamylated E1443. Individual b- and y-ion series and the amino acid sequence corresponding to each spectrum are indicated. Asterisks indicate ions with a neutral loss of a water molecule (see also **Supplementary Fig. 1**). (**c**) Michaelis-Menten fit of TTLL6 glutamylation of modified bovine brain or unmodified human microtubules (Methods). Error bars indicate s.e. of the fit (n=4).

Selectivity for the α-tubulin tail and the formation of long glutamate chains also holds when microtubules are used as substrate for TTLL6. LC-MS analyses of human unmodified microtubules glutamylated by TTLL6 show robust modification of the α- and not β-tubulin tail (**Supplementary Fig. 2a**). This substrate preference is consistent with cellular overexpression studies ^4^. MS-MS mapping of the TTLL6 glutamylation sites shows that the enzyme preferentially modifies E443 on the α-tail. Importantly, once the initial branch glutamate is added, our MS-MS analyses show that TTLL6 preferentially elongates the glutamate chain from this site as evidenced from the extracted ion chromatogram (**Supplementary Figs. 2b-k**). Consistent with it being an elongase, TTLL6 activity is low with unmodified microtubules and higher with brain microtubules, which are a heterogenous mix of mono- and polyglutamylated tubulin (**Fig. 1c**). The TTLL6 catalytic efficiency is 18-fold higher with glutamylated brain microtubules compared to unmodified microtubules, without a significant difference in the KM, indicating that glutamylation does not affect TTLL6 binding to microtubules, but affects significantly its ability to use the substrate for efficient glutamate transfer. Thus, TTLL6 has a strong preference for ligating glutamates to glutamates that have an exposed α-carboxyl group as opposed to internal glutamates in the tubulin tail that have only the γ-carboxyl group available for iso-peptide bond formation (Box 1). Consistent with this, we also find that TTLL6 has barely any activity on unmodified recombinant human αIA/βIII microtubules, but ~ 20-fold higher activity on detyrosinated αIA-Y/βIII recombinant human microtubules where the α-tubulin tail has a terminal glutamate with an available α-carboxyl group (**Supplementary Fig. 3**). Long glutamate chains have so far been detected only originating from internal glutamates in the tubulin tails, but the high activity of TTLL6 with detyrosinated microtubules raises the interesting possibility that long glutamate chains could also be added to the carboxy-terminus of α-tubulin *in vivo*.

### NMR shows glutamate chains catalyzed by TTLL6 are α-linked

Because our MS-MS analyses did not provide information on the connectivity of the glutamate chains, we used NMR to investigate whether TTLL6 adds α-linked (addition to the α-carboxyl group) or γ-linked (addition to the γ-carboxyl group) chains exclusively or generates chains with mixed linkages. Both types of linkages have been reported *in vivo* ^2,3^. Specifically, we used TTLL6 to incorporate ^13^C/^15^N-labeled glutamate into synthesized α-tail peptides with a pre-existing glutamate branch at position E445 (the position of the branch in the synthesized peptide was confirmed using 2D HH-TOCSY and NOESY experiments; **Methods** and **Supplementary Fig. 4**). Next, we used a ^1^H/^13^C-HSQC experiment to assign atom type to the chemical shifts of the added ^13^C/^15^N-Glu labeled residues by comparison to known chemical shifts for glutamate residues (Biological Magnetic Resonance Data Bank, **Supplementary Table 1**). ^1^H/^13^C-HSQC spectra contain peaks for any ^13^C-labeled atom directly bound to a proton and therefore contain resonances for all aliphatic atoms of the added glutamates. Peaks were further assigned as either ‘terminal’ (at the end of the added glutamate chain) or ‘intermediate’ (in the middle of the glutamate chain) by comparison of the HSQC data to a 2D CBCA(CO)(N)H spectrum (**Fig. 2a**). 2D CBCA(CO)(N)H spectra correlate any aliphatic carbon atom that is located two to three bonds from an amide (^1^H-^15^N) group with the proton of that amide, with the caveat that the magnetization must pass through a carbonyl ^13^C atom. If the added ^13^C/^15^N-glutamate residues have an α linkage, then the CBCA(CO)(N)H spectrum is expected to contain a C_α_ and C_β_ peak for each amide group that has a distinct chemical shift for the proton (**Figs. 2b** and **2c**). Conversely, if the linkage type is γ, the spectrum is expected to contain a C_α_ and C_β_ peak (**Figs. 2b** and **2d**). The presence of C_α_ and C_β_ peaks indicates that TTLL6 exclusively elongates α-linked chains (**Fig. 2**).

**Figure 2.**
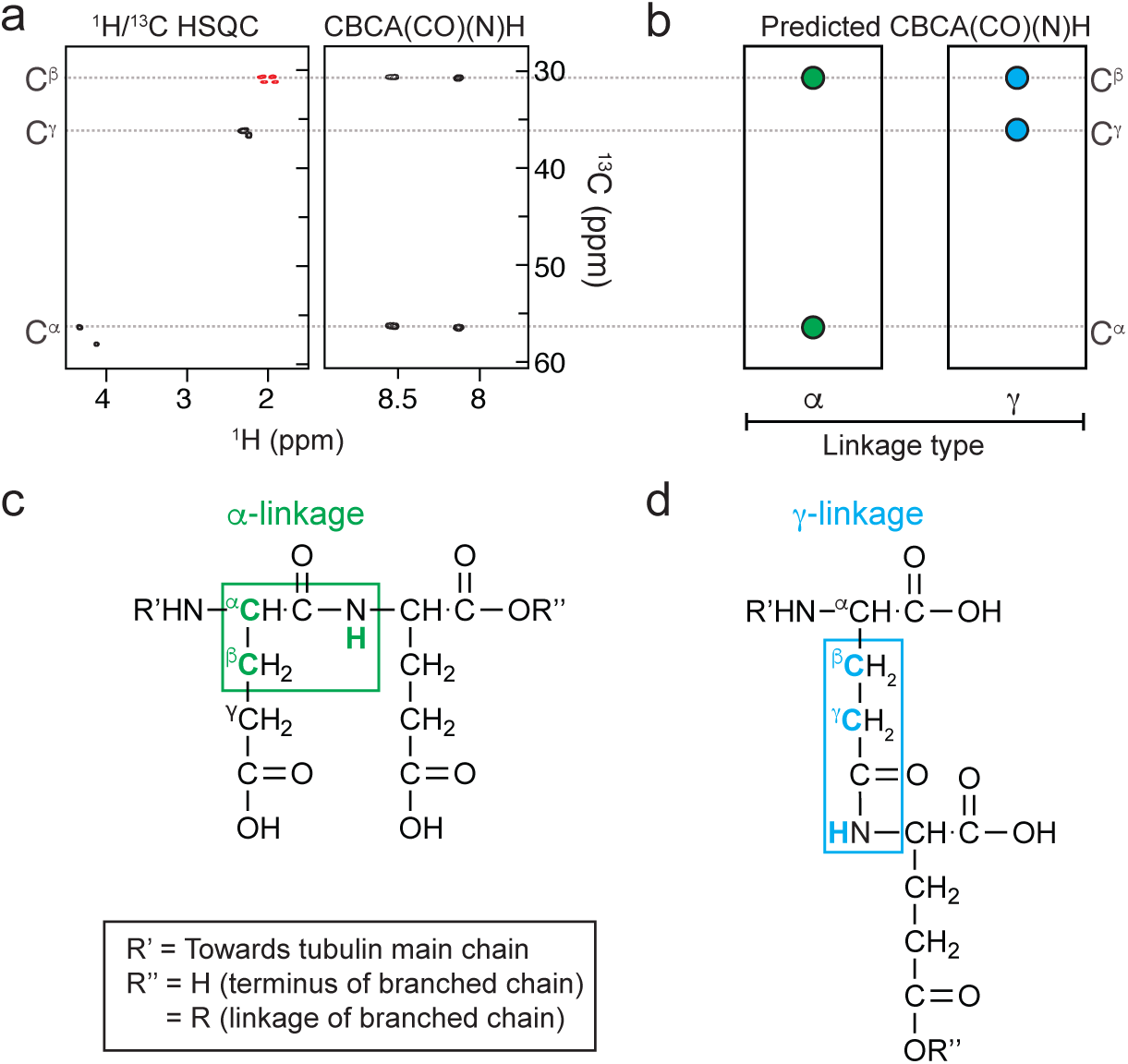
NMR spectroscopy indicates that enzymatic ^15^N/^13^C-glutamate addition occurs *via* an α linkage type. **(a)** A ^1^H/^13^C heteronuclear single quantum coherence (HSQC) spectrum was used to determine the aliphatic carbon and proton chemical shifts of the added ^15^N/^13^C-glutamates. A 2D CBCA(CO)(N)H experiment was used to identify the connectivity of the added glutamates. The spectrum showed that the two closest carbon atoms to amide carbonyls in the added glutamates belonged to C^α^ and C^β^ atoms, as predicted for an α-linkage type **(b)** prediction of the spectrum for an α- and γ-linkage type. **c)** In a CBCA(CO)(N)H experiment, magnetization originates in aliphatic protons and transfers *via* aliphatic carbons, through neighboring carbonyl and nitrogen amide atoms, before being observed on an amide proton (highlighted in green and blue boxes in **(c)** and **(d)**). Atoms that can be observed in the CBCA(CO)(N)H spectrum are highlighted in green **(c)** for the α-linkage type (C^α^, C^β^ and amide proton) and blue **(d)** for the γ linkage type (C^β^, C^γ^ and amide proton).

### TTLL6 is preferentially inhibited by a phosphinate α-elongation analog

Based on the structural conservation between tubulin amino acid ligases ^6,34,35^ and ATP-dependent ligases like D-Ala:D-Ala ligase ^36^, TTL and TTLLs are thought to employ a common catalytic mechanism that proceeds through phosphorylation of the carboxylate of an acceptor residue with the γ-phosphate of ATP to form an acyl-phosphate intermediate, followed by attack of the amino group of the incoming residue to be ligated to produce a tetrahedral intermediate that then eliminates the phosphate to yield a peptide bond (^36^; **Box 1**; **Supplementary Fig. 5**). Phosphinate analogs of this tetrahedral intermediate have been shown to be potent inhibitors for ATP-dependent amino acid ligases ^37^. In order to probe the substrate specificity and reaction mechanism of TTLL6, we synthesized three analogs in which a phosphinate replaces the tetrahedral carbon of the intermediate for the initiation, α-elongation or γ-elongation steps (**Fig. 3a**; ^38^). The analogs mimic a pseudo di-Glu peptide where the donor glutamic acid is linked either to the α- or γ-carboxylic acid of an acceptor glutamic acid through a phosphinate. In the case of the initiation analog, the main chain residues flanking the branch point glutamate are replaced by ethylamine (C-terminal side) and acetate (N-terminal side) (**Fig. 3a**).

**Figure 3.**
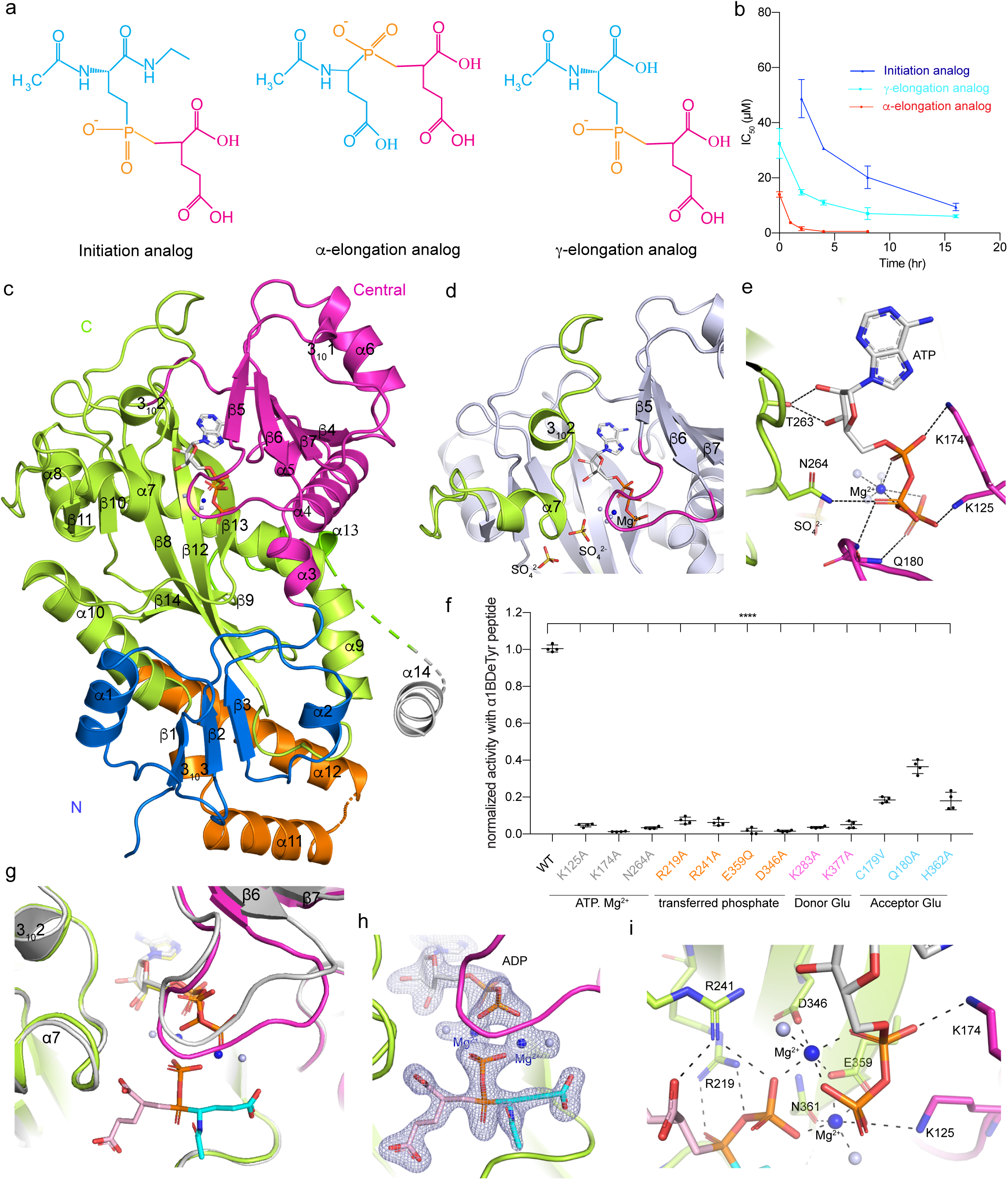
Structures of TTLL6 in complex with ATP and the α-elongation tetrahedral intermediate analog. (**a**) Chemical structures of the α-elongation analog, γ-elongation analog, and initiation analog of the glutamylation reaction; the acceptor and donor glutamate are shown in cyan and magenta, respectively. (**b**) IC_50_ values for the three analogs as a function of TTLL6 incubation time. Error bars indicate s.e. of the fit (n=4). (**c**) Ribbon diagram of the TTLL6 core bound to ATP. Nucleotide shown as a stick model. Dotted lines represent regions of the TTLL6 polypeptide chain that are disordered in the crystal structure. (**d**) Close-up view of the active site. (**e**) Close-up view showing residues important for ATP phosphate coordination. Nucleotide shown as a stick model. (**f**) Normalized *in vitro* glutamylation activity of recombinant structure guided TTLL6 mutants with the α1B-Y peptide. Error bars indicate s.e.m. (n=4). (**g**) Superposition of TTLL6 complexed with ATP (grey) and the α-elongation analog. (colored as in (**c**)), showing the displacement of the β5-β6 active site loop. Nucleotide and α-elongation analog shown as a stick model. The donor glutamate, transferred phosphate and acceptor glutamate of the α-elongation analog are colored pink, orange and cyan, respectively. Magnesium ion and water shown as blue and white sphere, respectively. (**h**) Active site showing the |*F*_o_|-|*F*_c_| density (prior to modeling the α-elongation analog) contoured at 5σ (grey). (**i**) Close-up of the active site showing residues important for γ-phosphate transfer and stabilization; hydrogen bonds denoted by dashed lines.

In the absence of any pre-incubation with ATP, we find that both the α- and γ-elongation analogs inhibit TTLL6 by more than 90%, with IC_50_ values of 13.2 μM and 33.9 μM, respectively. The initiation analog inhibits only ~ 20% of the enzyme’s activity, even at concentrations as high as 400 μM (**Fig. 3b** and **Supplementary Fig. 5d,** time = 0hr). Since this class of inhibitors is known to exhibit time-dependent inhibition ^37,39^, we pre-incubated the enzyme with ATP, MgCl_2_ and analogs before measuring activity. Inhibition for all three analogs increases markedly with pre-incubation time, with the α-elongation analog being the most potent (**Fig. 3b** and **Supplementary Fig. 5d**). The inhibition efficacy reaches saturation after ~ 4 hours of pre-incubation, with an IC_50_ value of ~ 600 nM (**Supplementary Fig. 5d**). The γ-linked elongation analog reaches saturation at ~ 8 hours with an IC_50_ value of ~ 6 μM (**Supplementary Fig. 5e**). The initiation analog requires the longest time to reach saturation (16 hours) with an IC_50_ value of 6.1 μM (**Supplementary Fig. 5f**). Taken together, these data show that TTLL6 has a strong preference for binding the α-elongation analog, consistent with our NMR findings that the enzyme catalyzes the formation of α-linked polyglutamate chains.

### Structures with tetrahedral intermediate analogs reveal molecular basis for α-linked glutamate chain elongation

To elucidate the structural features that specialize TTLL6 for catalyzing the formation of α-linked polyglutamate chains, we determined the crystal structures of the conserved catalytic core of mouse TTLL6 (res. 51-502) in complex with its ATP cofactor, as well as with the three tetrahedral intermediate analogs characterized above (**Table 1**, **Figs. 3c**, **4** and **Methods**). The TTLL6-ATP complex crystal structure was refined to 2.18 Å resolution with a *R*_free_ of 27.09%. Structures of the complexes with the three inhibitors were refined to 1.98 Å with a *R*free of 19%, 2.33 Å with a *R*_free_ of 21.25%, 2.51 Å with a *R*_free_ of 24.15%, for the α-elongation, γ-elongation and initiation analogs, respectively (**Table 1** and **Methods**). The TTLL6 core is elongated (~ 80 x 50 x 47 Å^3^) and consists of N-(res. 57-117), central (res. 118-206), and C-(res. 207-460) domains (**Fig. 3c**), similar to the closely related glutamylase TTLL7 ^6^, the glycylase TTLL3 ^40^ and the tubulin tyrosine ligase TTL ^34,35^. The ATP binds at the interface between the central and C-domains with the ribose and phosphate moieties cradled by two conserved loops, one in the central domain connecting strands β5 and β6 (res. 175-183), and one in the C-domain (res. 261-265) connecting helices 3102 and α7 (res. 261-265) that sit on top of the active site (**Fig. 3c, d**). Specifically, the sugar hydroxyls H-bond to T263, the α-phosphate is coordinated by K174, the β-phosphate by both N264 and the backbone amides of Q180 and G181, and the γ-phosphate interacts with the Q180 backbone amide and K125 in the N-domain (**Fig. 3e**). Both loops as well as helices 3_10_2 and α7 were not previously resolved in a TTLL7 structure in which the ATP β-phosphate had a high degree of disorder and the γ-phosphate was not visible (**Fig. 3d**;^6^), consistent with these loops undergoing a disorder-to-order transition upon productive binding of the ATP γ-phosphate. Consistent with their importance for nucleotide coordination, mutation of K174, K125, and N264 to alanine reduces TTLL6 activity to background levels (**Fig. 3f**).

**Figure 4.**
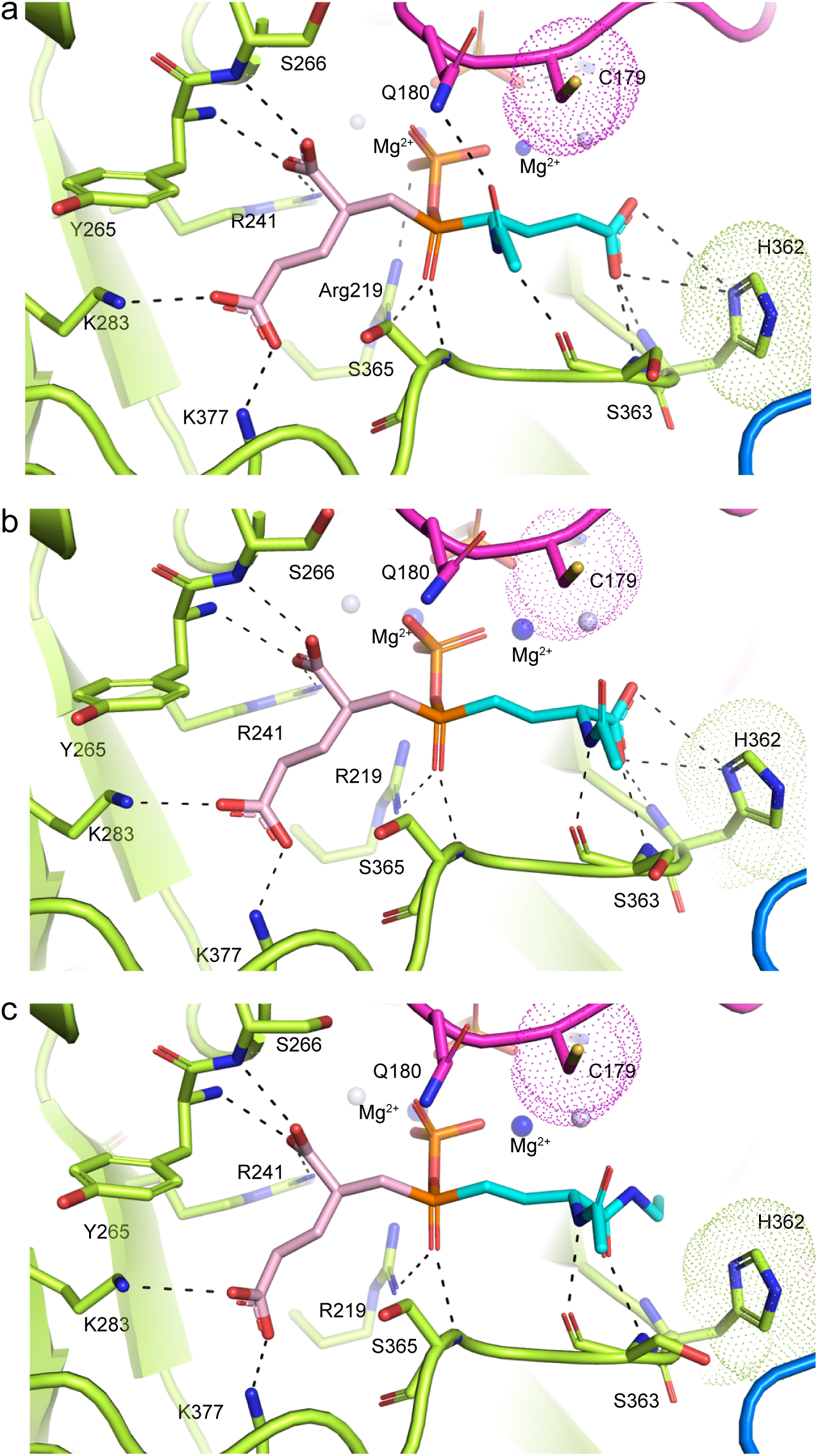
Structural basis for TTLL6 α-linked glutamate chain elongation activity. (**a-c**) Close-up views of the active site showing residues important for the recognition of the donor and acceptor glutamates in the α-elongation (**a**), γ-elongation (**b**) and initiation (**c**) tetrahedral intermediate complexes. Color scheme as in **Fig. 3g.**

**Table 1.**

| | ATP | α-elongation<br>analog complex | γ-elongation<br>analog complex | initiation<br>analog complex |
| --- | --- | --- | --- | --- |
| **Data collection** |  |  |  |  |
| Space group | P2<sub>1</sub> | P2<sub>1</sub> | P 2<sub>1</sub> | P 2<sub>1</sub> |
| Cell dimensions |  |  |  |  |
| *a*, *b*, *c* (Å) | 58.6, 115.7, 75.5 | 75.3, 110.6,<br>173.4 | 75.1, 109.7,<br>173.0 | 75.3, 110.6, 173.6 |
| $\alpha$, $\beta$, $\gamma$ (°) | 90.0,90.3,90.0 | 90.0, 90.3,90.0 | 90.0, 90.3,90.0 | 90.0,90.2,90.0 |
| Resolution (Å) | 45.93 - 2.18<br>(2.26 - 2.18) | 44.57 – 1.97<br>(2.048 – 1.98) | 44.29 – 2.33<br>(2.37 – 2.33) | 19.92-2.51<br>(2.594-2.50) |
| *R*<sub>merge</sub> | 0.021 (0.279) | 0.103 (0.495) | 0.092 (0.445) | 0.045(0.333) |
| *I* / σ*I* | 14.14 (2.49) | 16.23 (2.51) | 12.80 (2.12) | 9.95 (2.00) |
| Completeness (%) | 99.73 (98.91) | 97.69 (92.68) | 99.06 (95.22) | 98.10 (87.02) |
| Redundancy | 2.0(2.0) | 6.4(5.4) | 3.3(3.1) | 2.0(2.0) |
| **Refinement** |  |  |  |  |
| Resolution (Å) | 45.93 - 2.18 | 44.57 – 1.97 | 44.29 – 2.33 | 19.92-2.51 |
| No. reflections | 105,265 (10,321) | 193,618 (18263) | 118,734 (11355) | 194852 (19357) |
| *R*<sub>work</sub> / *R*<sub>free</sub> | 0.2372 / 0.2709 | 0.1669/ 0.1877 | 0.1821 / 0.2125 | 0.2091/ 0.2415 |
| No. atoms |  |  |  |  |
| Protein | 6173 | 13147 | 13042 | 12487 |
| Ligand/ion | 164 | 643 | 544 | 430 |
| Water | 141 | 993 | 593 | 287 |
| *B*-factors |  |  |  |  |
| Protein | 72.22 | 45.63 | 51.33 | 63.37 |
| Ligand/ion | 72.23 | 49.79 | 56.54 | 59.89 |
| Water | 63.20 | 48.57 | 49.01 | 49.77 |
| R.m.s. deviations |  |  |  |  |
| Bond lengths (Å) | 0.002 | 0.008 | 0.003 | 0.003 |
| Bond angles (°) | 0.53 | 0.96 | 0.58 | 0.61 |
| Co-ordinate error(Å) | 0.31 | 0.18 | 0.24 | 0.32 |

\*Values in parentheses are for highest-resolution shell.

**Table 1 (continued)**
| C179A/Q180R/H362I<br>Mutant | α-elongation<br>analog complex | γ-elongation<br>analog complex | initiation<br>analog complex |
| --- | --- | --- | --- |
| **Data collection** |  |  |  |
| Space group | P2<sub>1</sub> | P 2<sub>1</sub> | P 2<sub>1</sub> |
| Cell dimensions |  |  |  |
| *a*, *b*, *c* (Å) | 75.0, 109.5,<br>171.7 | 74.7, 108.6,<br>170.9 | 75.1, 108.8, 171.6 |
| $\alpha$, $\beta$, $\gamma$ (°) | 90,90.01,90 | 90,90.023,90 | 90,90.02,90 |
| Resolution (Å) | 46.16 – 3.08<br>(3.19 – 3.08) | 45.31 – 2.66<br>(2.75-2.66) | 45.95-2.60<br>(2.69-2.60) |
| *R*<sub>merge</sub> | 0.0605 (0.3552) | 0.04598 (0.3299) | 0.041 (0.312) |
| *I* / σ*I* | 7.16 (2.65) | 8.69 (2.02) | 10.45 (2.30) |
| Completeness (%) | 99.6 (98.36) | 99.64 (98.35) | 99.73 (99.07) |
| Redundancy | 2.0 (2.0) | 2.0 (2.0) | 2.0 (2.0) |
| **Refinement** |  |  |  |
| Resolution (Å) | 46.16 – 3.08 | 45.31 – 2.66 | 45.95-2.60 |
| No. reflections | 101188 (9901) | 155190 (15,060) | 84901 (8329) |
| *R*<sub>work</sub> / *R*<sub>free</sub> | 0.2190/ 0.2516 | 0.21362/ 0.2516 | 0.2271/ 0.2451 |
| No. atoms |  |  |  |
| Protein | 12480 | 12370 | 12691 |
| Ligand/ion | 346 | 458 | 418 |
| Water | 96 | 212 | 381 |
| *B*-factors |  |  |  |
| Protein | 89.7 | 68.54 | 59.07 |
| Ligand/ion | 69.86 | 63.62 | 56.22 |
| Water | 53.98 | 53.67 | 49.62 |
| R.m.s. deviations |  |  |  |
| Bond lengths (Å) | 0.002 | 0.002 | 0.002 |
| Bond angles (°) | 0.54 | 0.49 | 0.54 |
| Co-ordinate error(Å) | 0.35 | 0.37 | 0.33 |

\*Values in parentheses are for highest-resolution shell.

Binding of the α-elongation analog is accompanied by an inward movement of the β5-β6 loop involved in phosphate binding (maximal displacement ~ 5.7 Å; **Fig. 3g**). Interestingly, the β5-β6 loop was disordered in a TTL structure in complex with both ATP and the acceptor glutamate from the tubulin tail ^35^, suggesting that this structural element becomes ordered when bound to the tetrahedral state intermediate. Close inspection of the TTLL6 active-site electron density reveals that the γ-phosphate of the ATP was transferred to the phosphinate moiety in the analog to form ADP and a phosphorylated inhibitor, with continuous density clearly visible from the analog to the phosphoryl group (**Fig. 3h**). As a result, the ADP β-phosphate breaks its interaction with the Gln180 backbone amide and makes additional interactions with K125, while the γ-phosphate that was transferred to the analog breaks its interaction with K125 and makes new interactions with R219 and R241. Consistent with their importance in the coordination of the transferred γ-phosphate, mutation of R219 or R241 reduces TTLL6 activity to background levels (**Fig. 3f**). The equivalent residues in TTL, R202 and R222, are also essential for activity ^34^, consistent with a common catalytic mechanism. The phosphoryl group is linked to the β-phosphate of the ADP through two Mg^2+^ cations. These are coordinated by N346 and E359 (**Fig. 3i)** and neutralize the charge repulsion between the two highly negatively charged phosphates. Mutation of D346 and E359 also reduces TTLL6 activity to background (**Fig. 3f**). The phosphorylated inhibitor more closely mimics the true tetrahedral intermediate (which also bears a phosphate) than the unmodified phosphinate inhibitor (**Supplementary Fig. 5**). This phosphate transfer occurs in all three inhibitor complexes (**Fig. 4**, **Supplementary Figs. 6a, 6b, 6d**) and explains the time-dependent inhibition of TTLL6 by these analogs. Phosphate transfer to the analogs is the rate-limiting step of the overall inhibition process, and the enzyme has higher affinity for the phosphorylated analogs. Similar observations were made for D-Ala:D-Ala ligase ^36^ that catalyzes the ligation of D-Ala to D-Ala in an ATP-dependent manner, supporting a shared enzymatic mechanism.

The donor glutamate is nestled in a highly positively charged groove (**Supplementary Figs. 6d**) and is coordinated the same way in all three analog complex structures. The α-carboxylate of the donor glutamate H-bonds to the backbone amides of Y265 and S266 and the guanidinium of invariant R241 (**Figs. 4a-c**). The γ-carboxylate of the donor glutamate is stabilized by electrostatic interactions with K283 and K377 (**Figs. 4a-c**). The side chain of an aspartate would be too short for these interactions, explaining why it cannot be used as a substrate by TTLL glutamylases. In the ATP complex, a sulfate ion is bound at the site occupied by the γ-carboxylate in the analog complex structures (**Fig. 3d**). These two lysines are invariant across all TTLL glutamylases but not so in glycylases, which ligate a glycine to a glutamate, or TTL which ligates a tyrosine to a glutamate (**Supplementary Fig. 7**). All glycylases have either hydrophobic or polar side chains at these positions, suitable for glycine coordination ^40^. All TTLs have a glutamate in place of K238, suitable for H-bonding, and a tyrosine in place of K377, suitable for π stacking interactions with the donor tyrosine ^34,35^. Mutation of either K283 or K377 to alanine in TTLL6 reduces its activity to background levels (**Fig. 3f**).

The major differences between the α- and γ-linked elongation analogs complex structures are in the interactions made by the acceptor glutamate of the analog with the enzyme. In the case of the α-linked elongation analog, the carbonyl and amide groups of the peptide bond N-terminal to the acceptor glutamate H-bond to the side chain of invariant Q180 and the backbone carbonyl oxygen of S363, respectively (**Fig. 4a**). Mutation of Q180 to alanine decreases TTLL6 activity by 64% (**Fig. 3f**). In the case of the γ-linked elongation analog, the carbonyl group of the peptide bond N-terminal to the acceptor glutamate is no longer within hydrogen bonding distance to Q180 (**Fig. 4b, c**), explaining the less potent inhibition (**Fig. 3b**). The non-reactive carboxylate of the acceptor glutamate in both complexes (the γ-carboxylate for the α-linked elongation analog and the α-carboxylate for the γ-linked elongation analog) is coordinated in the same manner through a H-bond with the backbone amide of S363 and a salt bridge with H362 (**Figs. 4a, b**). Mutation of H362 to alanine reduces TTLL6 activity by 82% (**Fig. 3f**). In the case of the initiation analog, the salt bridge with H362 is lost, consistent with the less potent inhibition by this analog (**Fig. 3b**). A positively charged groove runs adjacent to the acceptor glutamate (**Supplementary Fig. 6d**) that is optimal for binding a negatively charged peptide such as the tubulin tail in the early stages of the glutamylation reaction or the growing polyglutamate chain.

### Active site molecular determinants for glutamate chain elongation *versus* initiation

Our TTLL6 structures in complex with the three tetrahedral intermediate analogs point to Q180 in the β5-β6 loop (**Fig. 4a**) as one of the key residues for specifying whether a TTLL enzyme has a preference to elongate or initiate glutamate chains. The conservation patterns of this residue among TTLLs glutamylases suggests a segregation between initiases and elongases with TTLL1, 6, 9, 11, and 13 as elongases, and TTLL2, 4, 5 and 12 as initiases (**Supplementary Fig. 7a**). The residue equivalent to TTLL6 Q180 is an arginine in all human TTLL glutamylases thought to be initiases based on cellular overexpression studies and antibody detection (^4^; **Supplementary Fig. 7a**). To test this classification, we purified recombinant TTLL4 core (res. 106-601; **Methods**) and examined its activity with unmodified tubulin tail peptides and tubulin tail peptides with branched glutamates. Unlike, TTLL6, TTLL4 prefers to glutamylate the β-tubulin tail over the α-tail (**Fig. 5a**). Importantly, and in contrast to TTLL6, a detyrosinated α-tubulin peptide with a glutamate at the carboxy-terminus or an α-tail peptide with a branch glutamate does not increase TTLL4 activity (**Fig. 5a**). We mapped the sites TTLL4 glutamylates on the branched βI^442^E_1_ peptide and found that the enzyme adds glutamates to E439 and E440, but leaves the branched glutamate untouched (**Figs. 5b, 5c** and **Supplementary Fig. 8a**). Thus, TTLL4 catalyzes the formation of an iso-peptide bond with the γ-carboxylate of a glutamate in the tubulin tail to initiate a branch. This specificity for catalyzing the addition of a branch glutamate on the β-tubulin also holds when microtubules are used as substrates. LC-MS analyses of microtubules glutamylated by TTLL4 indicates that the enzyme prefers to modify β-tubulin (**Supplementary Fig. 8b**) and MS-MS demonstrates that it adds only single glutamates on the β-tubulin tails at multiple positions (**Supplementary Figs. 8c-f**). We detected up to three posttranslationally added glutamates on β-tubulin (**Supplementary Fig. 8b**), with all of them added as mono-glutamates at multiple internal glutamates on the β-tubulin tail (**Supplementary Figs. 8c-f**). Consistent with TTLL4 being an initiase, the *k*_cat_ of the enzyme is 4-fold higher with unmodified microtubules compared to heterogenous brain microtubules which are glutamylated, but also contain unbranched glutamates that could still serve as substrates for TTLL4 (**Fig. 5d**). The *K_M_* is similar between the two substrates, indicating that substrate glutamylation does not significantly affect TTLL4 binding and only its ability to catalyze glutamate ligation. Thus, TTLL4 and TTLL6 function as glutamate chain initiase and elongase, respectively, and while the binding energies derived from the interaction with the entire tubulin substrate are similar regardless of whether it is glutamylated or not, the discrimination arises during catalysis proper.

**Figure 5.**
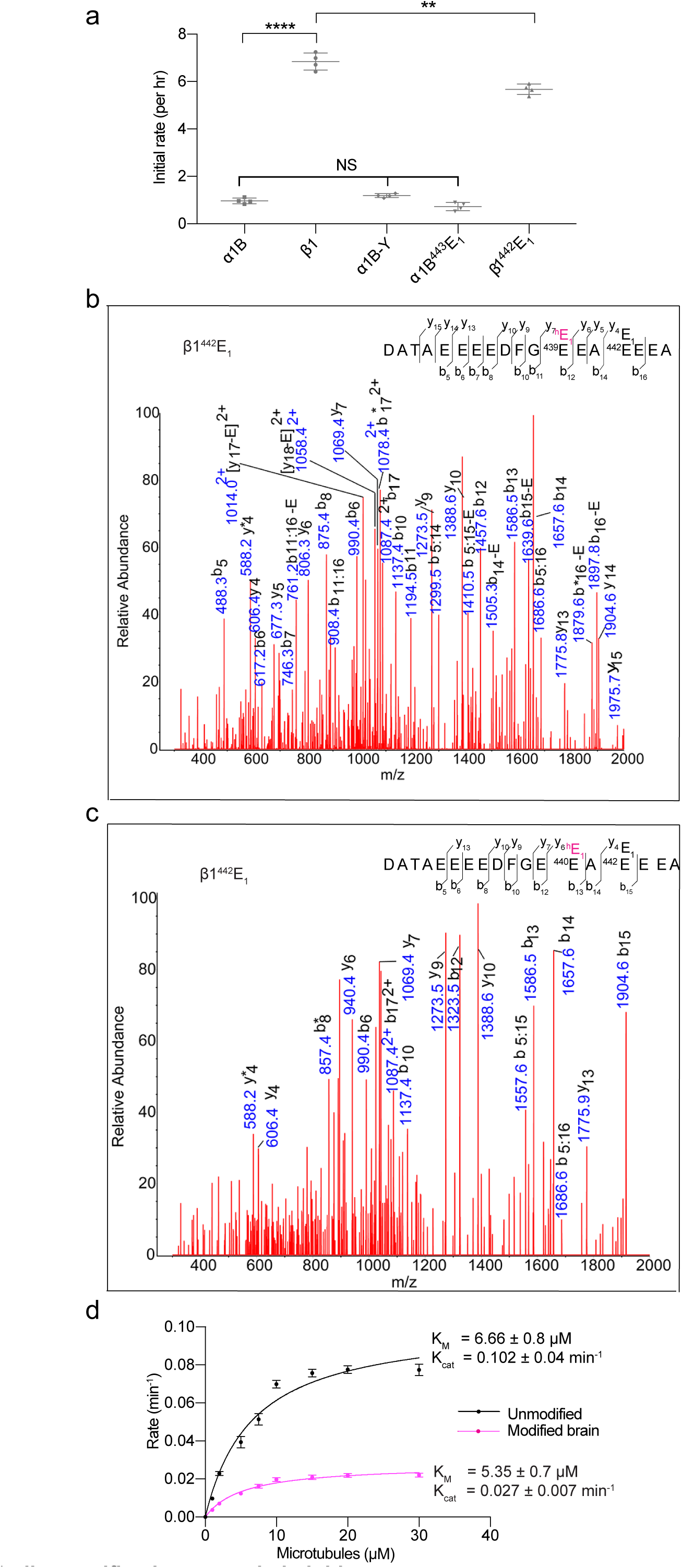
TTLL4 is a β-tubulin specific glutamate chain initiase. (**a**) Glutamylation of α- and β-tubulin tail peptides by TTLL4. Error bars indicate s.e.m. (n=4). (**b, c**) MS/MS sequencing of β1^442^E_1_ peptide glutamylated by TTLL4. Enzymatically added monoglutamates are added at E439 (**b**) and E440 in (**c**). Individual b-, y-ion series and the amino acid sequence corresponding to each spectrum are indicated. Asterisks indicate ions with a neutral loss of a water molecule. (**d**) Michaelis-Menten fit of glutamylation of modified brain or unmodified human tubulin by TTLL4 (STAR Methods). Error bars indicate s.e.m of the fit (n=4).

### Molecular engineering switches TTLL6 preference from elongation to initiation

We then sought to further test our hypothesis by converting TTLL6 into an initiase by mutating into arginine Q180, which segregates between initiases and elongases. In addition, in comparing TTLL6 and TTLL4 active site residue conservation, two additional residues stood out as being potentially important for specifying initiation *versus* elongation. First, H362 which as an Ile in TTLL4 would relieve the buried charge of the histidine that would lack a salt bridge partner (the carboxylate of the acceptor glutamate) in the case of the initiation reaction and thus would be unfavorable. Second, C179 which when mutated to alanine could potentially relieve the steric clash with the carbonyl of the peptide bond N-terminal to the acceptor glutamate (**Fig. 4** and **Supplementary Fig. 7b**). Indeed, we find that this C179A/Q180R/H362I mutant TTLL6 no longer discriminates between heterogenous glutamylated brain microtubules and unmodified microtubules, as both the *k*_cat_ and *K*_M_ are similar with these substrates (**Fig. 6a**). This is in contrast to wild-type TTLL6 which shows an 18-fold enhancement in *k*_cat_ with glutamylated brain microtubules (**Fig. 1c**). Consistent with it now being an initiase that prefers to catalyze iso-peptide bond formation, the triple-mutant TTLL6 shows a strong preference for glutamylating an unmodified α1B peptide. While wild-type TTLL6 showed 15-fold higher elongation activity on a glutamylated peptide with a branched glutamate (1B^443^E_1_), the mutant enzyme no longer shows such strong discrimination (**Fig. 6b**). Moreover, mapping of the sites glutamylated by the C179A/Q180R/H362I TTLL6 mutant shows that unlike the wild type-enzyme, the mutant does not prefer to elongate from one site (**Supplementary Fig. 2b-k**). Instead, the dominant doubly glutamate species (~ 84.8%) show addition of mono-glutamates at multiple positions in the α-tubulin tail, consistent with this mutant functioning predominantly as an initiase (**Figs. 6c, d** and **Supplementary Fig. 9a, b**). While we were able to detect as many as four additional glutamates on the α-tubulin tails, extracted ion chromatograms indicate that only a small fraction of the tetra- and tri-glutamylated species had preferential elongation from E443 in the α-tail (16% and 9% of the wild-type elongation activity, respectively), with the majority being a mixture of glutamate additions at multiple positions in the tail (**Fig. 6e**). Consistent with this, the C179A/Q180R/H362I TTLL6 mutant is inhibited an order of magnitude less efficiently by the α-linked elongation analog compared to wild-type (IC_50_ of 5.4 ± 0.8 *versus* 0.6 ± 0.05 μM) and all three analogs now inhibit the mutant with comparable potency (**Fig. 6f**) with IC_50_ values of 3.5 ± 0.2 μM and 2.5 ± 0.1 μM for the initiation and γ-linked elongation analog, respectively. Mutation of two additional residues to more closely resemble an initiase active site did not further improve initiation activity of a pentamutant C179A/Q180R/R182I/H362I/S367H nor the inhibition by the initiation analog (**Fig. 6b** and **Supplementary Fig. 10a**). Taken together, our kinetic and mass spectrometry analyses coupled with inhibitor studies delineate three residues which interact with the acceptor glutamate that are important for the functional specialization of TTLL6 into an elongase.

**Figure 6.**
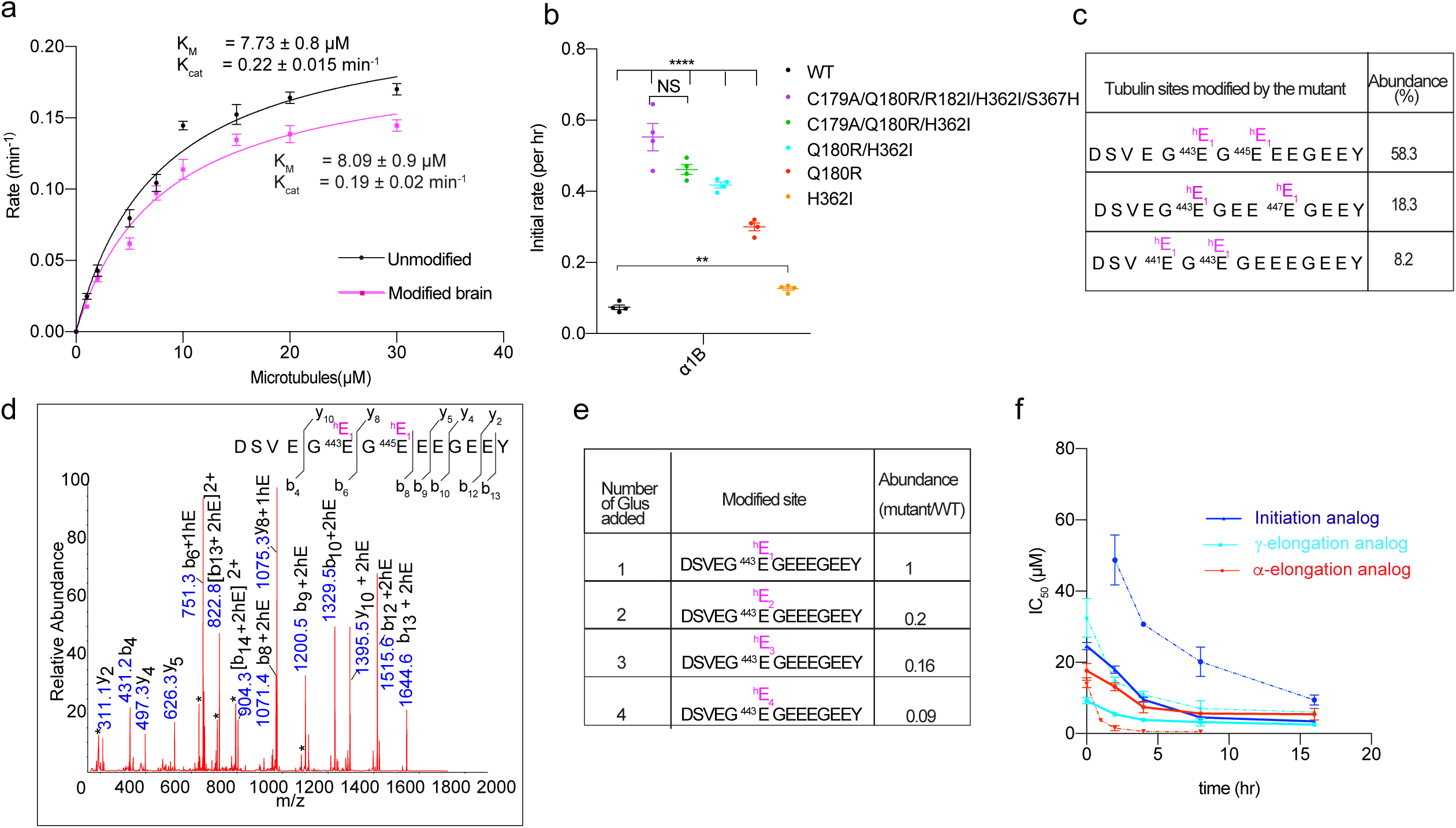
Structure-based engineering of TTLL6. (**a**) Michaelis-Menten fit for glutamylation of modified porcine brain (magenta) and unmodified human tubulin (black) by the TTLL6 C179A/Q180R/H362I mutant. (**b**) Activity of structure-based mutants with unmodified (α1B) synthesized α-tubulin peptide. Error bars shown in (**a, b**) indicate s.e.m. (n=4). (c) Table showing the most abundant di-glutamylated species glutamylated by the TTLL6 C179A/Q180R/H362I mutant (**d**) MS-MS sequencing of the di-glutamylated α-tubulin C-terminal tails of microtubules glutamylated by the TTLL6 C179A/Q180R/H362I mutant. Individual b-, y-ion series and the amino acid sequence corresponding to each spectrum are indicated. Asterisks indicate ions with a neutral loss of a water molecule. (**e**) Table showing the relative abundance detected for elongated glutamate chain species for the C179A/Q180R/H362I mutant and wildtype TTLL6. (**f**) IC_50_ values for the three tetrahedral intermediate analogs plotted as a function of incubation time for the TTLL6 C179A/Q180R/H362I mutant. IC_50_ values for the wildtype enzyme shown as dotted lines for comparison. Error bars indicate s.e.m of the fit (n=4).

### Structures of engineered mutants reveal mechanism of regioselectivity switch

To understand the structural basis for this change in selectivity, we solved X-ray crystal structures of the C179A/Q180R/H362I TTLL6 mutant in complex with the three analogs (**Table 1, Supplementary Fig. 10b-d** and **Methods**). We find that in the complexes with the initiation and γ-linked analogs the mutation of Q180 to arginine favors binding to these analogs because the side chain of an arginine at this position is long enough to H-bond with the carbonyl of the acceptor glutamate of the initiation and γ-linked elongation analogs (**Figs. 7a** and **b**). In the wild-type TTLL6, the Q180 side chain is too short to make this stabilizing interaction (5.7Å *versus* 3.2 Å; **Figs. 4a** and **b**). The glycine elongase TTLL10 also has an invariant glutamine at this position, while the two glycine initiases TTLL3 and 8, both have an invariant arginine (**Supplementary Fig. 6a**), indicating that this position in the active site is used to discriminate between α-peptide versus iso-peptide formation throughout the entire TTLL family and not only glutamylases. Mutation of H362 to isoleucine allows van der Waals stabilizing interaction with the ethyl in the initiation analog and also opens up the active site to accommodate the C-terminal tail of tubulin (**Figs. 7a** and **4c**), but fails to neutralize the buried charge of the acceptor carboxylate for the α and γ-linked elongation analogs (**Figs. 7b, c**). Consistent with this, a H362I mutant is 1.8-fold more active glutamylating an unmodified α-tubulin peptide than wild-type TTLL6 (**Fig. 6b)**. Interestingly, among all TTLLs that are elongases, only TTLL13 has a histidine at this site. However, all other elongating TTLLs have side chains in the vicinity that are able to reach and neutralize the α-carboxylate (H68 for TTLL1, R352 for TTLL7, N363 for TTLL9 and N442 for TTLL11). C179 is not critical in this discrimination, as a double Q180R/H362I mutant is almost as effective in initiating branch glutamates on the unmodified tubulin tail peptide as the C179A/Q180R/H362I triple mutant (**Fig. 6b**). In conclusion, the active site of TTLL6 has evolved for efficient glutamate ligation to the α-carboxylate of the acceptor glutamate, while the TTLL4 active site has evolved for efficient glutamate ligation to the γ-carboxylate of the acceptor glutamate. Two conserved residues in TTLL6, Q180 and H362, are key to the functional segregation between these initiase and elongase activities (**Fig. 6**).

**Figure 7.**
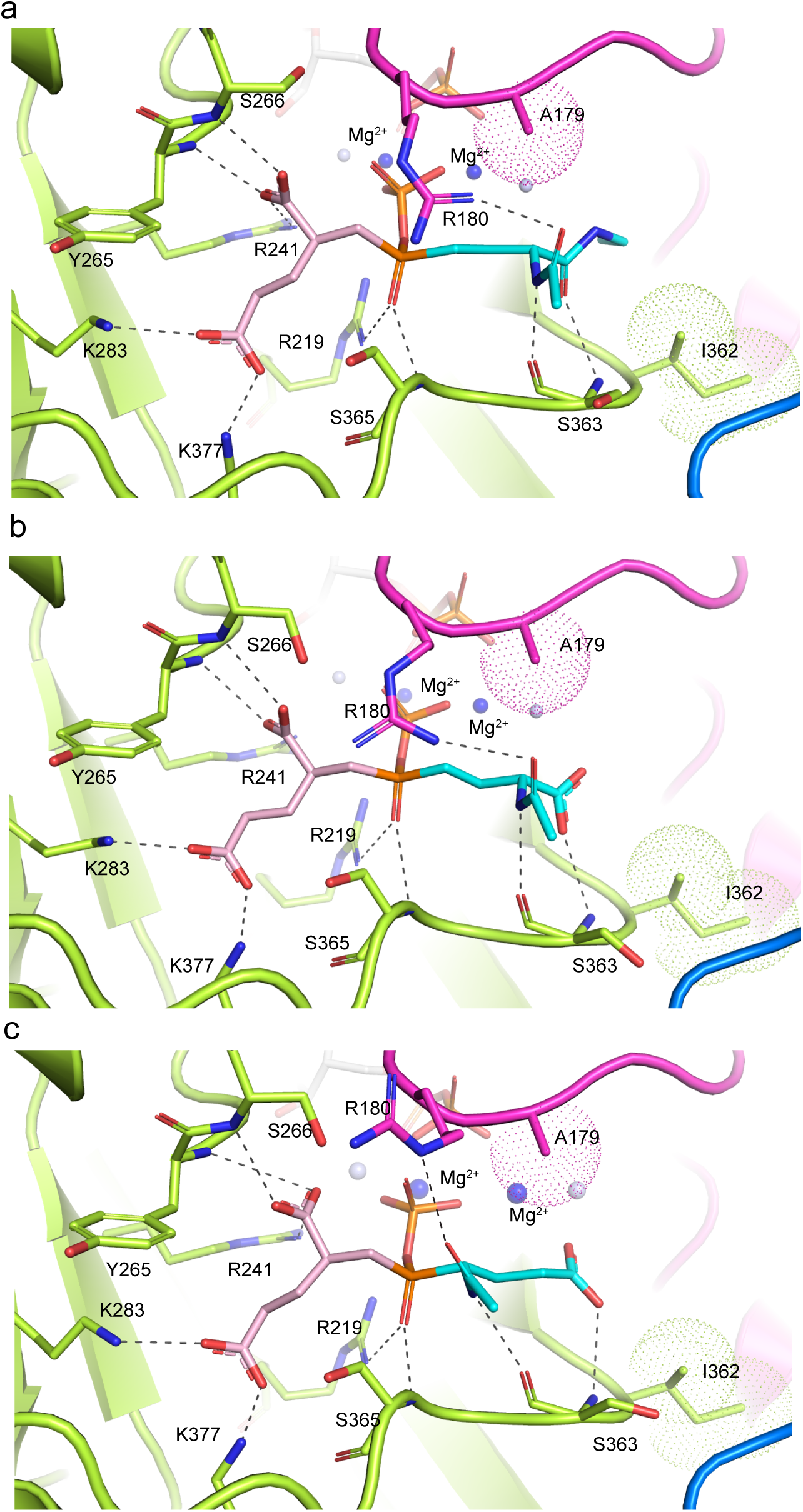
Structure engineered TTLL6 reveals basis for elongation *versus* initiation activity. (a-c) Close-up view of the TTLL6 C179A/Q180R/H362I mutant active site showing residues important for donor and acceptor glutamate recognition for the initiation (a), γ-elongation (b) and α-elongation (c) analogs.

## Discussion

Our combined functional, MS-MS and NMR analyses show that TTLL6 preferentially elongates α-linked glutamate chains (**Figs. 1** and **2**) and its activity increases with the number of glutamates in the extended glutamate branch, consistent with strong elongation activity. In contrast, TTLL4 does not elongate glutamate chains, but introduces mono-glutamates at multiple internal positions in the β-tubulin tail through a γ-linked or iso-peptide bond (**Fig. 5** and **Supplementary Fig. 8**). We report the first examples of time-dependent potent inhibitors of the TTLL family enzymes. Our high-resolution structures of TTLL6 in complex with ATP and three such inhibitors that mimic the initiation and α- and γ-linked elongation tetrahedral intermediates revealed the key active site residues that control this regioselectivity (**Figs. 3** and **4**) and that when mutated can switch TTLL6 to behave primarily as an initiase (**Fig. 6**). We show that among these Q180 is the main determinant for this regioselectivity. This residue segregates with initiases and elongases not only for TTLL glutamylases, but the entire TTLL family encompassing both glutamylases and glycylases. Thus, our structural and functional work lays bare the molecular basis for regioselectivity for all TTLL enzymes. Among amino acid ligases that use a glutamate as an acceptor, γ-linked reactions are most commonly observed in all kingdoms of life ^41^. The residues that are key for specifying α-linked elongase activity are invariant among TTLL6 orthologs from ciliates to humans, suggesting a strong selective pressure for the formation of long α-linked glutamate chains in eukaryotes.

The evolution and conservation of TTLL glutamylases with distinct regioselectivities suggests that readers of the tubulin code may be sensitive to the nature of the linkage structure of these glutamate chains. Previous studies have shown that microtubule associated proteins respond differently to short versus long polyglutamate chains. For example, tau, MAP1B and MAP2 bind to microtubules that are moderately glutamylated (1-3 glutamates; ^42–44^), while MAP1A specifically recognizes long glutamate chains (~ 7 glutamates; ^42^). Kinesin 1 and 2 also have higher processivity on α-glutamylated microtubules, with kinesin 1 showing increased processivity with a longer glutamate chain on the α-tail ^14^. Varying glutamate chain lengths on the β-tubulin tail provides graded regulation of microtubule severing ^16^. Both α- and γ-linked chains have been reported in adult brain tubulin ^3 2^. Thus, our study provides impetus for understanding how TTLLs with distinct regioselectivities are used by the cell to functionalize microtubules.

In addition to providing the first biophysical evidence for segregated glutamate chain initiation and elongation activities for TTLL enzymes, we show that TTLL4 and 6 discriminate between the α- and β-tubulin tails. This is in contrast to the glutamylase TTLL7 which does not discriminate between the isolated α- and β-tails ^6,7^ and preferentially glutamylates the β-tubulin tail only in the context of the microtubule lattice because of interactions through domains outside the enzyme catalytic core ^6^. A subsequent molecular modeling study concluded that TTLL glutamylase active sites in general cannot discriminate between α- and the β-tubulin tails ^45^ and proposed that tubulin tail choice is dictated exclusively by elements outside the catalytic core for all TTLL family glutamylases ^45^. Our experimental data demonstrate that, in fact, TTLLs have evolved different stringencies for substrate recognition by their active sites and that their strategies for tubulin recognition cannot be easily generalized without careful quantitative biochemical work. TTLLs broadly, and TTLL4 and TTLL6 in particular, have been recently linked to the glutamylation of multiple non-tubulin substrates which highlights the importance of understanding the active site proximal and distal determinants for the substrate specificity of this highly diversified enzyme family.

A major limitation towards our mechanistic understanding of the tubulin code has been the inability to generate microtubules with complex posttranslational modification patterns for *in vitro* reconstitution. By identifying the sites and patterns of modifications introduced by TTLLs, we not only provide insight into the biochemical program that generates the complex *in vivo* microtubule modification patterns, but also a tool for generating differentially modified microtubules for further analyses of the tubulin code. Furthermore, progress in elucidating how the tubulin code regulates cell physiology has been hampered by the inability to rapidly inhibit or activate tubulin modification enzymes and a lack of tubulin modification reporters for live cell imaging. Ours is the first structure of a glutamylase or any TTLL enzyme with an ordered active site, and in complex with its substrates, ATP, acceptor and donor amino acids. Enzymes in the ATP-grasp superfamily to which TTLL enzymes belong are known to use non-protogenic amino acids as substrates ^46^. Our structures enable the future rational engineering of TTLL active sites to be compatible with amino acid analogs, similar to the compensatory mutations made for kinases ^47^ that have revolutionized studies into their function. These analogs could also be functionalized for fluorescence or used for click chemistry for imaging in cells. Last, but not least, our structures will serve as a springboard for the design of inhibitors for therapeutic intervention in neurodegenerative disorders characterized by tubulin hyper-glutamylation or modulation of the innate immune response by cGAS glutamylation.

## Acknowledgements

We thank D-Y. Lee from the Biophysics Core (National Heart, Lung and Blood Institute) for access and advice for mass spectrometry. N.T. is supported by the intramural program of the National Heart, Lung and Blood Institute (NHLBI). A.R-M. is supported by the intramural programs of the National Institute of Neurological Disorders and Stroke (NINDS) and NHLBI. M.E.T. is supported by the Natural Sciences and Engineering Research Council of Canada (NSERC).

## Author Contributions

K.K.M. obtained crystals, collected X-ray data, solved and refined all structures, performed functional assays and mass spectrometry analyses. E.K.K. purified proteins, obtained the wildtype ATP and elongation inhibitor crystals, collected and processed X-ray data. M.S. collected and interpreted NMR data. Y. Li collected and interpreted MS-MS data. Y. Liu synthesized inhibitors. H.L.B. synthesized branched peptides. M.E.T. supervised Y. Liu. N.T. interpreted NMR data. A.R-M. initiated, coordinated and supervised project. K.K.M interpreted functional data. K.K.M and A.R.M. analyzed structures and wrote the manuscript with contributions from M.S. and N.T. All authors read and approved the manuscript.

## Accession Numbers

Atomic models have been deposited at the Protein Data Bank under accession numbers: PDB ID***, PDB ID***, PDB ID***, PDB ID*** for the wild-type TTLL6 complexed with ATP, α-elongation, initiation and γ-elongation analog, respectively and PDB ID***, PDB ID***, PDB ID*** for the TTLL6 mutant complexed with the α-elongation, initiation and γ-elongation analog, respectively.

## METHODS

### Protein expression and purification

*Mus musculus* TTLL6 (51-502) was expressed using *E. coli* Arctic Express DE3 cells (Agilent) with a GST tag on its N-terminus. Cultures were grown at 37 °C to an OD600 of 2. Protein expression was induced using 0.5 mM IPTG at 13 °C for 24 hours. Cells were lysed using a microfluidizer and cell lysate was clarified by centrifugation at 16000 rpm for 45 minutes. The fusion protein was purified using GST affinity chromatography and the GST tag was removed using 3C Prescission protease at 1:10 molar ratio. TTLL6 was further purified on a heparin column followed by size exclusion chromatography. Site directed mutagenesis was performed using Quickchange (Agilent) and all mutants were purified as the wildtype construct. All constructs eluted as a single peak from a Superdex 75 gel filtration column (GE). Mass spectrometry was used to analyze the purity and integrity of the wild-type as well as mutant constructs. All measured masses were found to be consistent with the theoretical mass within 0.1-1 Dalton.

*Caenorhabditis elegans* TTLL4 (106-601) was expressed using *E. coli* Arctic Express DE3 cells (Agilent) with a GST tag on its N-terminus. Cultures were grown at 37 ° C to an OD600 of 2. Protein expression was induced using 0.5 mM IPTG at 13 °C for 24 hours. Cells were lysed and the clarified cell lysate was purified using GST affinity chromatography. The GST tag was removed using 3C Prescission protease at 1:10 molar ratio. TTLL4 was further purified on a heparin column followed by size exclusion chromatography using a Superdex75 column (GE). The protein eluted as a single peak.

### Peptide Synthesis

Peptides were synthesized on either low load Wang resins (Novabiochem) or super acid sensitive resin for protected peptides, using an automated Applied Biosystems 433A synthesizer. Amino acids were purchased from P3Bio or Novabiochem, except H-Glu(tBu)-OtBu, Fmoc-Glu(2-Phenylisopropyl ester)-OH which were purchased from Bachem. Novasyn TG Sieber resin was purchased from Novabiochem. The backbone peptide sequence was synthesized with orthogonally protected Fmoc-Glu(2-Phenylisopropyl ester)-OH at the branch point. Fully protected Glu polypeptides were synthesized on super acid sensitive Novasyn TG Sieber resin and cleaved using 1% TFA in DCM. The combined 1% TFA washes were combined,, neutralized with DIEA, and rotovaped under reduced pressure. The orthogonally protected Glu was deprotected with 1% TFA in DCM. The protected polypeptide Glu chains were coupled to the resin-bound peptide using PyBOP or PyOxim coupling reagents overnight. Final cleavage of the branched peptides was achieved using 95% TFA containing scavengers 1,2-ethandithiol and thioanisole (1:2). The cleavage reaction was performed at room temperature for 1.5-3h depending on the length of the peptide. The cleaved peptide was precipitated in diethyl ether, centrifuged and washed several times with fresh ether. Purification of the crude peptides was performed on a C18 Grace-Vydac semi-preparative reversed-phase HPLC column (250×10mm) using a Waters 600 series HPLC system. Fractions were analyzed on a Vydac C18 analytical column (150×4.6mm). Separations were achieved using linear gradients of 0-100% buffer B for 120 mins (semipreparative) or 30 mins (analytical), at a flow rate of 3ml/min or 1ml/min, respectively. Buffer A was water containing 0.045% TFA and buffer B was acetonitrile with 0.036% TFA. Detection was achieved at 220nm. The identity of the peptides was confirmed using MALDI-TOF on a Waters LR mass spectrometry instrument.

### Peptide glutamylation assays

Synthetic peptides were dissolved in dH2O and the pH was adjusted to 7.0 using KOH. Glutamylation reactions of the peptides were performed by mixing 250μL of solution containing 20 mM Hepes pH 7.0, 50 mM KCl, 10 mM Mg Cl_2_, 5 mM TCEP, 1 mM ATP, 1 mM L-glutamic acid, 20 μM peptide and 1 μM TTLL6 or 100 nM TTLL4. The reaction was initiated by the addition of the enzyme. Reaction samples were collected at 0, 2, 4, 6, 8, 16, 24, and 30 hours. Reactions were quenched by the addition of an equal volume of 20% Acetonitrile with 0.1% TFA and separated on a Zorbax 300SB C18 column (Agilent) using a 0-70% acetonitrile gradient in 0.05% TFA at a flowrate of 0.2 mL/min. The column was attached in line with a 6224 ESI-TOF LC/MS (Agilent) and the data were analyzed using the Mass Hunter Workstation platform as previously described ^6,40^. Activity assays of TTLL6 structure guided mutants were performed with the α1B-Y peptide at a 1:10 enzyme to substrate molar ratio (2.5 μM enzyme and 25 μM peptide) in the same buffer condition as above. Reaction samples were collected at 0, 2, 4, 6, 8, 16, 24, and 30 hrs. The decrease in the m/z signal of the unmodified peptide was monitored as a function of time. The enzyme itself was used as an internal loading standard, as it elutes separately from the tubulin peptides. Reactions were performed in duplicate on separate days.

### Glutamylation assays

Glutamylation activity of wildtype and mutant TTLL6 constructs was determined by quantifying the incorporation of radio-labeled [^3^H]-glutamate into 20 μM taxol stabilized brain microtubules. The reactions were carried out in buffer containing 20 mM HEPES, pH 7.0, 10 mM MgCl_2_, 10 mM KCl, 1 mM DTT, 1 mM ATP, 20 uM Taxol, 95 μM ^2^H-glutamate and 5 μM ^3^H-glutamate (49.6 Ci/mmol). Reactions were initiated at 30° C by the addition of the enzyme to a final concentration of 1 μM. Aliquots were removed at different time points and reactions were terminated by the addition of 20 mM EDTA. The reaction mix was spotted on Amersham Hybond-N^+^ membrane (GE Lifesciences). Tubulin binds to the filter and unincorporated radioactivity was removed by washing with a phosphate buffer containing 25 mM KCl. The radioactivity associated with the filters was then quantified using a scintillation counter and the total glutamate transferred to tubulin was calculated from the total radioactivity in the reaction mix. Reactions were performed in duplicate on two different days (n=4). Initial rates were measured by fitting the linear range of the curve using the program package PRISM 6.

The three intermediate state analogs in which a phosphinate replaces the tetrahedral carbon of the intermediate for the initiation, α-elongation or γ-elongation steps were synthesized as described previously^38^. The initiation and γ-elongation analogs are a mixture of two diastereomers and the α-elongation analog is a mixture of four. The three inhibitor IC_50_ values for the three tetrahedral intermediate analogs were determined by varying the concentration of the analogs and measuring the amount of tritium labelled L-Glutamic acid transferred to taxol-stabilized brain microtubules by TTLL6. The assays were performed at 30°C. The reaction mix consists of 20 mM Hepes, pH 7.0, 50 mM KCl, 10 mM MgCl_2_, 1 mM TCEP, 1 mM ATP, 950 μM L-Glutamic acid, 50 μM H^3^-L-Glutamic acid, and 10 μM brain microtubules. The analogs were examined for their ability to stimulate a time-dependent inhibition of the enzyme by pre-incubating 1 μM TTLL6, 1 mM ATP in buffer solution containing 20 mM HEPES pH 7.0, 50 mM KCl and 10 mM MgCl_2_ with a varying concentration of the analogs. Aliquots were withdrawn at 0, 1, 2, 4, 8, and 16 hours and added to the reaction mix. Reactions were allowed to proceed for the indicated times and terminated by the addition of saturating amounts of EDTA and incubation on ice for 30 minutes. The reaction mix was then transferred to the positively charged nylon transfer membrane (GE Healthcare) and washed extensively with 50 mM sodium phosphate buffer, pH 7.8 and 25 mM KCl. The residual radioactivity was measured using a Beckman scintillation counter. IC_50_ values were obtained by fitting the data to the four parameter logistic curves in Prism 6.1.

Taxol stabilized microtubules were polymerized from recombinant α1A/βIII and α1A-Y/βIII tubulins and incubated with TTLL6 at a 1:10 molar ratio (enzyme:tubulin) in 20 mM Hepes pH 7.0, 50 mM KCl, 10 mM Mg Cl_2_, 5 mM TCEP, 1 mM ATP, 1 mM L-Glutamic acid, 20 μM taxol and 1 μM TTLL6. Aliquots were removed at the indicated time points and the reaction quenched by the addition of EDTA to a final concentration of 50 mM followed by addition of SDS loading buffer. Five μg of protein was loaded per lane, separated on a 8-12% gradient gel and transferred on to a nitrocellulose membrane. Mouse monoclonal antibody GT335 (AG-20B-0020, Adipogen) was used to detect tubulin glutamylation. IRDye700 Goat anti-mouse secondary antibody was used for primary antibody detection. The GT335 signal was normalized to total tubulin levels using rabbit monoclonal anti-α-tubulin antibody (EP1332Y, Abcam). Densitometric analyses and quantifications were done using the Image Studio Lite software package (LI-COR Biosciences).

### Kinetic analyses of microtubule glutamylation

The K_M_ and k_cat_ values for TTLL6 wild-type, the switch mutant (C179A/Q180R/H362I) and TTLL4 with brain and unmodified microtubules were determined by varying the concentration of tubulin from 1 to 20 μM for TTLL6 and 1 to 30 μM for TTLL4 while keeping the enzyme concentration constant at 1 μM. The reaction condition consisted of 50 mM Hepes pH 7.0, 10 mM MgCl_2_, 50 mM KCl, 1 mM DTT, 1 mM ATP, 950 μM L-Glutamic acid, and 50 μM H^3^-L-Glutamic acid. Reactions were initiated by the addition of the enzyme and terminated by the addition of saturating amounts of EDTA and placing it on ice for 30 mins. The reaction mix was then transferred to the positively charged nylon transfer membrane (GE Lifesciences) and washed extensively with 50 mM sodium phosphate, pH 7.8 supplemented with 25 mM KCl. The membranes were then transferred to a scintillation vial and residual radioactivity was measured using a Beckmann scintillation counter. Kinetic constants were obtained by using the one site binding parameter equation in Prism 6.1. Reactions were performed in duplicate on two separate days.

### NMR sample preparation and analyses

TTLL6 was incubated with the branched peptide α1B^445^E_1_ at room temperature in a reaction mix containing 20 mM Hepes, pH 7.0, 50 mM KCl, 10 mM MgCl_2_, 1 mM TCEP, 1 mM ATP, 1 mM ^15^N L-glutamic acid (Sigma Aldrich, 332143) or ^13^C5 ^15^N L-glutamic acid (Sigma Aldrich, 607851), and 200 μM peptide. The reaction was initiated by the addition of 20 μM TTLL6 enzyme and quenched by the addition of equal volume of 20% acetonitrile with 0.1% TFA after 16 hr incubation at room temperature. The modified peptide was then separated from the reaction mix on a Grace Vydac C18 column (Fisher Scientific, 218tp54) using a 0-70% acetonitrile gradient in 0.05% TFA at a flow rate of 0.5 mL/min. Fractions containing the peptide were collected, lyophilized and then dissolved in a buffer solution containing 50 mM sodium phosphate, pH 6.8 and 50 mM NaCl.

D_2_O was added to the ^15^N Glu- or ^13^C/^15^N Glu-modified α1B^445^E_1_ peptide to a final concentration of 7% and the sample placed in a Shigemi tube (typical sample volume of 250 μL). In both samples, the α1B^445^E_1_ peptide and the first added Glu were unlabeled, while the rest of the enzymatically added Glu residues were isotopically labeled. ^1^H/^13^C-CT-HSQC (CT = 1/*J*_CC_) and 2D CBCA(CO)(N)H NMR data were acquired for the ^15^N/^13^C Glu-labeled sample at 298 K on a Bruker 600 MHz spectrometer equipped with a cryogenic probe. The 2D CBCA(CO)(N)H experiment was adapted from a standard 3D CBCA(CO)NH by omitting ^15^N chemical shift evolution. Experiments for the ^15^N Glu-labeled sample included a 2D HH-TOCSY measured at 600 MHz and a 2D HH-NOESY (90 ms mixing time) measured at 800 MHz, both with a cryoprobe. Spectra were processed using NMRPipe and peak-picked using NMRDraw ^48^.

### Crystallization and structure determination for wild-type TTLL6 complexes

All crystals were obtained by hanging drop vapor diffusion at 16 ° C. For the TTLL6-ATP-Glu co-crystals, 7.5-8.0 mg/mL of the protein solution containing 1 mM ATP and 1 mM L-glutamic acid was mixed with an equal volume of crystallization reservoir solution comprised of 0.2 M ammonium sulfate and 10-15 % of PEG 3350. Small crystals appeared overnight and grew to a maximum size (~300 x 200 x 40 μm) within a week. Crystals grew with symmetry P21 with 2 copies per asymmetric unit (Unit cell: *<u>a</u>*=58.6 Å, *<u>b</u>*=75.5 Å, *<u>c</u>*=115.7 Å, α=γ=90°, β=90.3°; diffraction limit=2.18 Å). Crystals of TTLL6 complexed with the α-elongation, γ-elongation and initiation analogs were obtained by mixing 8 mg/mL of TTLL6 containing 1 mM ATP and 1 mM of the inhibitor at 1:1 ratio with 100 mM sodium citrate, pH 6.2, 200 mM MgCl_2_ and 8-12 % PEG 20K. Crystals grew to maximum dimensions of ~ 800 x 300 x 50 μm within 7-10 days with symmetry P21 with 4 copies per asymmetric unit (Unit cell for TTL6-α-elongation analog: *<u>a</u>*=75.3 Å, *<u>b</u>*=110.6 Å, *<u>c</u>*=173.4 Å, β=90.2°; diffraction limit=1.977 Å; Unit cell for TTL6-γ-elongation analog: *<u>a</u>*=75.1 Å, *<u>b</u>*=109.7 Å, *<u>c</u>*=173.0 Å, β=90.3°; diffraction limit=2.33Å; Unit cell for TTL6-initiation analog: *<u>a</u>*=75.2 Å, *<u>b</u>*=110.6 Å, *<u>c</u>*=173.6 Å, β=90.1°; diffraction limit=2.5 Å). Prior to data collection, all crystals were cryo-protected by quickly soaking in fresh reservoir solution containing 30 % glycerol. Crystals were flash frozen by directly plunging into liquid nitrogen. X-ray diffraction data for ATP bound TTLL6 were collected at the Advanced light source (ALS) beamline 5.0.1 while the data sets for the inhibitor complexes were collected at ALS beamlines 5.0.2 and 5.0.3. The diffraction data for the ATP complex and the initiation analog complex were indexed and integrated using the XDS program suite ^49^. The integrated data were scaled using Scala in the CCP4 program suite ^50^. The diffraction data for the α-elongation analog and the γ-elongation analog complexes were indexed, integrated and scaled using HKL2000 ^51^.

The structures complexed with the three analogs were solved by molecular replacement using PHASER-MR ^52^ as implemented in Phenix ^53^ using the TTLL7 structure ^6^ (PDB ID:4YLR) as the search model without the microtubule binding domain. Residues around the active site were excluded from the search model. Difference electron density maps revealed unambiguous density for the microtubule binding domain (residues 392-459) and the ATP. Several rounds of iterative model building and refinement were performed using COOT^54^ and Phenix, respectively. The current refined model of TTLL6-ATP includes residues 57 - 460, 2 ATP nucleotides, 141 water molecules, 2 magnesium and 8 sulfate ions. Residues 410-416 of the polypeptide chain B were not well-resolved in the electron density map and are presumed disordered. The density for the C-terminal helix was not of sufficient quality to assign register unambiguously and was left as a polyalanine model. The Rwork and Rfree of the TTLL6-ATP model are 23.72 and 27.09 %, respectively (**Table 1**). MolProbity revealed no unfavorable (ϕ,Ψ) combinations, and the structural parameters of both main and side chains were found to be better than average.

The current refined model of the ATP-α-elongation analog complex structure consists of residues 57-459 for all four copies in the AU (with the exception of copy C for which residues 407-415 had poorly resolved electron density and were left unmodeled), 4 phosphorylated analogs, 943 water molecules, 58 glycerol molecules, 8 magnesium and 2 chlorine ions. The density for the C-terminal helix was not of sufficient quality to assign register unambiguously and was left as a poly-alanine model. The ligand was built and its molecular geometry was calculated by eLBOW^55^ as implemented in Phenix. The electron density for the phosphorylated inhibitor is consistent with a L-glutamate stereoisomer. The Rwork and Rfree of the current TTLL6-ATP model are 16.69 and 18.77 %, respectively (**Table 1**). MolProbity revealed no unfavorable (ϕ,Ψ) combinations, and the structural parameters of both main and side chains were found to be better than average.

The current refined model of the ATP-γ-elongation analog structure consists of residues 57-459 for all four copies in the AU, 4 phosphorylated analogs, 593 water molecules, 8 magnesium ions, 38 glycerol molecules. The Rwork and Rfree of the current TTLL6-ATP model are 18.21 and 21.25 %, respectively (**Table 1**). MolProbity revealed no unfavorable (ϕ,Ψ) combinations, and the structural parameters of both main and side chains were found to be better than average.

The current refined model of the ATP-initiation analog structure consists of residues 57-459 for all four copies in the AU (with the exception of residues 330-333 of chain A, residues 412-417 of chain B and residues 411-414 of chain C which were poorly resolved in the electron density and left unmodeled), 4 phosphorylated analogs, 287 water molecules, 8 magnesium ions and 33 glycerol molecules. The R_work_ and R_free_ of the current TTLL6-ATP model are 20.91 and 24.15%, respectively (**Table 1**). MolProbity revealed no unfavorable (ϕ,Ψ) combinations, and the structural parameters of both main and side chains were found to be better than average.

### Crystallization and structure determination for mutant TTLL6 complexes

Crystals for the C179A/Q180R/H362I TTLL6 mutant in complex with the γ-elongation and initiation analogs were obtained under similar conditions as for the wild-type construct. The complex with the α-elongation analog also crystallized in similar conditions, but 2mM of the compound was used in the crystallization drop. Crystals grew to maximum dimensions of ~ 800 x 300 x 50 μm within 7-10 days with the symmetry P2_1_ with 4 copies per asymmetric unit (Unit cell for TTLL6 C179A/Q180R/H362I-α-elongation analog: *<u>a</u>* 75.03 Å, *<u>b</u>*=109.49 Å, *<u>c</u>*=171.7 Å, α=γ=90°, β=90.014°; diffraction limit=3.084 Å; Unit cell for TTL6 C179A/Q180R/H362I-γ-elongation analog: *<u>a</u>*=74.73 Å, *<u>b</u>*=108.64 Å, *<u>c</u>*=170.88 Å, α=γ=90°, β=90.023°; diffraction limit=2.65Å; Unit cell for TTL6-C179A/Q180R/H362I initiation analog: *<u>a</u>* 75.099 Å, *<u>b</u>*=108.81 Å, *<u>c</u>*=171.61 Å, α=γ=90°, β=90.025°; diffraction limit=2.6 Å). Crystal cryoprotection, data collection and reduction were performed as for the wild-type crystals. All mutant structures were solved by molecular replacement with the TTLL6 structure with the ATP and all residues in the active site removed. The current refined model of the TTLL6 C179A/Q180R/H362I mutant in complex with ATP and the α-elongation analog consists of residues 57-459 for all four copies in the AU (with the exception of copies A and B for which residues 413-416 and residues 412-416, respectively had poorly resolved electron density and were left unmodeled), 4 phosphorylated analogs, 96 water molecules, 18 glycerol molecules, 8 magnesium ions. The Rwork and Rfree of the current model of TTLL6 C179A/Q180R/H362I mutant in complex with the α-elongation analog are 21.90% and 25.16 %, respectively (**Table 1**). MolProbity revealed no unfavorable (ϕ,Ψ) combinations, and the structural parameters of both main and side chains were found to be better than average.

The current refined model of the TTLL6 C179A/Q180R/H362I mutant in complex with ATP and γ-elongation analog consists of residues 57-459 for all four copies in the AU (with the exception of copy B for which residues 411-413 had poorly resolved electron density and was left unmodeled), 4 phosphorylated analogs, 212 water molecules, 39 glycerol molecules, 8 magnesium ions. The Rwork and Rfree of the current model are 21.62% and 25.16%, respectively (**Table 1**). MolProbity revealed no unfavorable (ϕ,Ψ) combinations, and the structural parameters of both main and side chains were found to be better than average.

The current refined model of the TTLL6 C179A/Q180R/H362I mutant in complex with ATP and initiation analog consists of residues 57-459 for all four copies in the AU (with the exception of copies A and B for which residues 412-415 and residues 412-413, respectively had poorly resolved electron density and were left unmodeled), 4 phosphorylated analogs, 377 water molecules, 28 glycerol molecules, 8 magnesium ions. The Rwork and Rfree of the current TTLL6-ATP model are 22.71 % and 24.51%, respectively (**Table 1**). MolProbity revealed no unfavorable (ϕ,Ψ) combinations, and the structural parameters of both main and side chains were found to be better than average.

### Mass spectrometric analysis of intact glutamylated tubulin

Unmodified tubulin was purified from tsA201 cells using a TOG affinity column as described in ^56,57^. Taxol stabilized unmodified microtubules were incubated with TTLL6 or TTLL4 at a 1:10 molar ratio (enzyme:tubulin) in 20 mM Hepes pH 7.0, 50 mM KCl, 10 mM Mg Cl_2_, 5 mM TCEP, 1 mM ATP, 1 mM L-glutamic acid, 20 μM taxol and 1 μM TTLL6 or TTLL4. Aliquots were removed at the indicated time points. The reaction was quenched by the addition of an equal volume of 0.1% trifluoroacetic acid, 20 mM EDTA, and 10% acetonitrile. Samples were analyzed by electrospray mass spectrometry as described in ^16^. The spectra display the characteristic distribution of masses with peaks separated by +129 Da corresponding to one glutamate.

### MS-MS analyses of glutamylated microtubules and peptides

Microtubules were polymerized by mixing 1:10 v/v of DMSO:tubulin at 37° C for 1hr and subsequently stabilized by the addition of 20 μM taxol. Any unpolymerized tubulin was separated by passing it through a 60% glycerol cushion in 1xBRB80 buffer (80 mM Pipes, 1 mM MgCl_2_, 1 mM EGTA and pH 6.8) through centrifugation at 37 ° C for 15 min at 65000 rpm. The pellet was then resuspended in 1xBRB80 buffer containing 20 μM taxol and 1 mM DTT. TTLL6 was incubated with taxol-stabilized unmodified microtubules at room temperature (1 μM TTLL6, 10 μM microtubules) in buffer solution consisting of 20 mM Hepes pH 7.0, 50 mM KCl, 10 mM MgCl_2_, 5 mM TCEP, 1 mM ATP, 1 mM L-glutamic acid, and 20 μM taxol, and aliquots removed at 0h, 15 min, 1, 2, 4 and 8 hrs. Reactions were quenched by the addition of an equal volume of SDS-PAGE loading buffer. 3 μg of glutamylated microtubules were separated by SDS-PAGE as described ^58^. Bands for α-tubulin were excised, and digested with Asp-N (1:20 ratio of protease:substrate) at 37 °C overnight. Digests were extracted from the gel and desalted with Waters Oasis HLB μElution plate prior to LC/MS/MS analysis. A Thermo-Fisher Orbitrap Elite mass spectrometer coupled with a 3000 Ultimate HPLC was used for LC/MS/MS data acquisition. Samples were separated with Mobile Phase A (MPA): 0.1% formic acid and Mobile Phase B (MPB): acetonitrile with 0.1% formic acid and gradient: 0 min, 2% MPB, 30 min, 27% MPB, 90 min, 120 min, 80%MPB. The Thermo-Fisher Scientific Orbitrap Elite mass spectrometer was operated in positive nano-electrospray mode. Unless otherwise indicated, the resolution of the survey scan was set at 60k at m/z 400 with a target value of 1 x 10^6^ ions. The m/z range for MS scans was 300–1600. The MS/MS data were acquired in data-dependent mode with an isolation window of 2.0 Da; Dynamic exclusion of 18s. Unless otherwise indicated, minimal signal required for triggering MS-MS data acquisition is 1 x 10^4^ ions. The top fifteen most abundant ions were selected for product ion analysis. Modifications related to ^13^C glutamylation were custom-built and searched against all spectra corresponded to glutamylated peptides were manually curated.

Peptides were modified as described in the previous section and subjected to LC-MS-MS analysis. A Thermo-Fisher Orbitrap Elite mass spectrometer coupled with a 3000 Ultimate HPLC was used for all LC-MS-MS data acquisition. For the α1B and α1BDeTyr peptides, samples were separated on a gradient: 0 min, 2% MPB, 66 min, 30% MPB, 90 min, 80% MPB. The top twenty most abundant ions were selected for product ion analysis. For the α1B^443^E_1_ and α1B^445^E_1_ peptides, samples were separated on a gradient: 0 min, 2% MPB, 32 min, 27% MPB, 60 min, 80% MPB. The top ten most abundant ions were selected for product ion analysis. The MS-MS data were acquired in data-dependent mode with an isolation window of 1.5 Da and the minimal signal required for trigging MS-MS data acquisition is 1 x 10^3^. For the β1^442^E_1_ peptide, samples were separated on a gradient: 0 min, 2% MPB, 32 min, 36% MPB, 60 min, 80% MPB. The top six most abundant ions were selected for product ion analysis. The MS-MS data were acquired in data-dependent mode with an isolation window of 1.5 Da. Modifications related to ^13^C glutamylation were custom-built and searched against all spectra corresponded to glutamylated peptides were manually curated.

