## Supplementary Table and Figures for "Structural basis for polyglutamate chain initiation and elongation by TTLL family enzymes"

**Supplementary Table 1: Chemical shift assignments for the $^{13}C/^{15}N$ TTLL6 glutamylated $\alpha 1B^{445}E_1$ peptide.**

| Atom | Intermediate $\delta$ (ppm) | Terminus $\delta$ (ppm) | BMRB Glu $\delta$ ±S.D. (ppm) |
|---|---|---|---|
| $H\alpha$ | 4.331 | 4.123 | 4.238 ± 0.401 |
| $H\beta 2$ | 2.078 | 2.045 | 2.021 ± 0.207 |
| $H\beta 3$ | 1.943 | 1.912 | 1.997 ± 0.213 |
| $H\gamma 2$ | 2.305 | 2.237 | 2.268 ± 0.209 |
| $H\gamma 3$ | 2.305 | 2.237 | 2.249 ± 0.213 |
| $C\alpha$ | 56.239 | 57.966 | 57.330 ± 2.065 |
| $H_N$ | 8.522 | 8.129 | 8.330 ± 0.580 |
| $C\beta$ | 30.700 | 31.222 | 29.973 ± 1.698 |
| $C\gamma$ | 36.218 | 36.654 | 36.103 ± 1.205 |

Atom assignments were made by comparison to protein glutamate chemical shifts derived from the Biological Magnetic Resonance Data Bank (filtered to remove proteins bound to paramagnetic or aromatic ligands, dated 22nd May 2018). Assignment to intermediate or terminal glutamate was determined using the CBCA(CO)(N)H experiment.

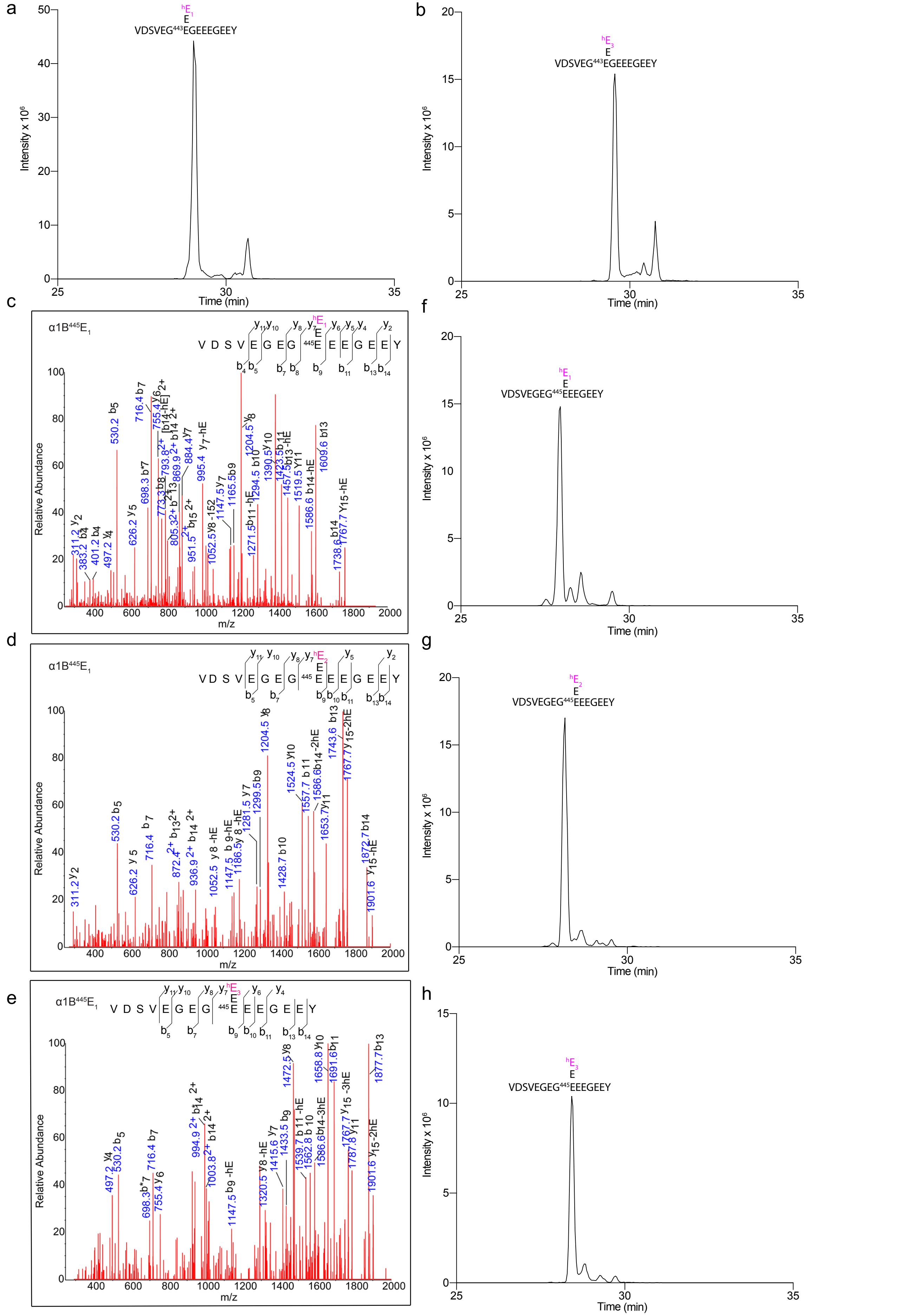


**Supplementary Figure 1**. **TTLL6 is an α-tubulin elongase.**
(a, b) Extracted-ion chromatogram of TTLL6 modified α1B$^{443}$E$_1$ peptide to which one (a) and three heavy glutamates (b) were added to the existing mono-glutamate at position E443. (c-e) MS/MS sequencing of α1B$^{445}$E$_1$ peptideglutamylated by TTLL6 showing the addition of one (c), two (d) and three (e) heavy glutamates to the existing mono-glutamate at positionE445. Individual b- and y- ion series and the amino acid sequence corresponding to each spectrum are indicated. Asterisks indicate ions with a neutral loss of a water molecule.
(f-h) of TTLL6 modified α1B$^{445}$E$_1$ peptide to which one (f), two (g) and three heavy glutamates (h) were added to the existing mono-glutamate at position E445.

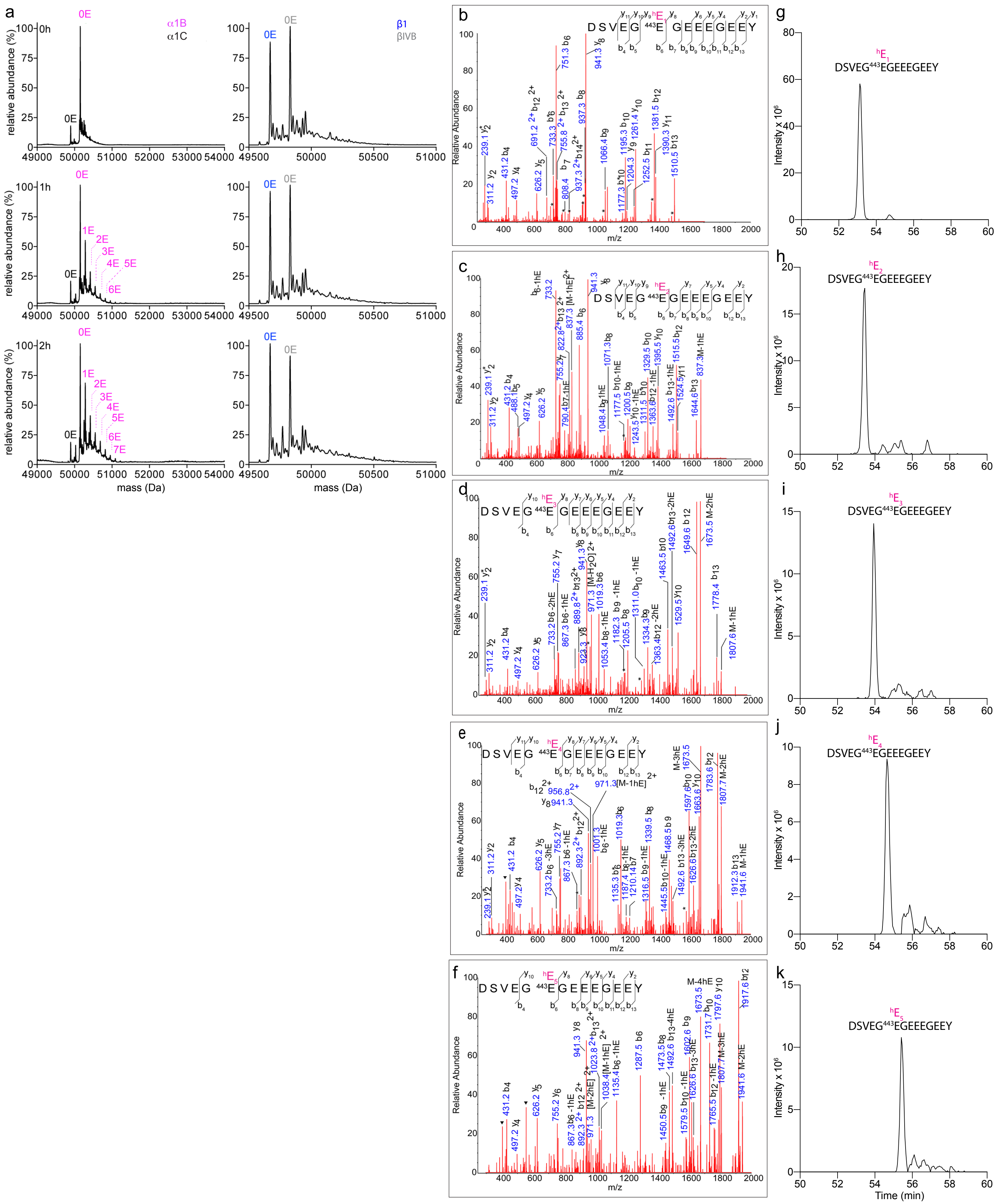


**Supplementary Figure 2. TTLL6 is an α-tubulin specific elongase.**
(a) Deconvoluted α- and β-tubulin mass spectra of unmodified human microtubules glutamylated by TTLL6 (STAR Methods). The number of added glutamates is indicated and colored according to tubulin isoform. (b-f) MS/MS sequencing of the α-tubulin C-terminal tails of microtubules glutamylated by TTLL6. Mono-glutamylated species in (b), di-glutamylated species in (c), tri-glutamylated species in (d), tetra-glutamylated species in (e), and penta-glutamylated species in (f). Individual b-, y-ion series and the amino acid sequence corresponding to each spectrum are indicated. Asterisks indicate ions with a neutral loss of a water molecule. (g-k) Extracted-ion chromatogram of mono-glutamylated (g), di-glutamylated (h), tri-glutamylated (i), tetra-glutamylated (j) and penta-glutamylated (k) C-terminal α-tubulin tail proteolytically excised from microtubules showing modification at E443.

a

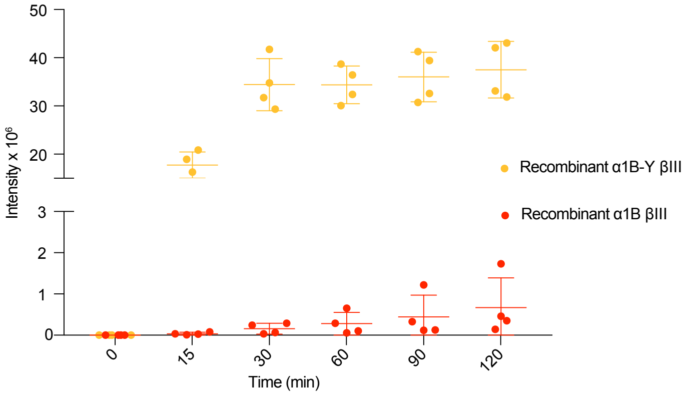


**Supplementary Figure 3. TTLL6 preferentially glutamylates detyrosinated α-tubulin in recombinant human microtubules.** TTLL6 glutamylation activity with recombinant α1B-Y/βIII (orange) and α1B/βIII (red) microtubules. Error bars indicate s.e.m (n=4).

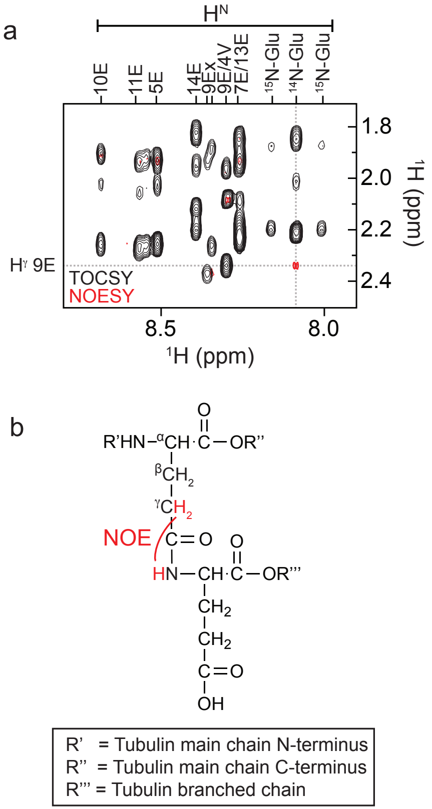


**Supplementary Figure 4. NMR spectroscopy confirms that the synthetic peptide used as a TTLL6 substrate has a glutamate branch at E445.**

(a) A zoom region of the overlaid 2D HH-TOCSY and NOESY experiments. The 2D TOCSY correlates proton spin systems. The region selected shows the $H^N$ (x-axis) and glutamate $H^\beta$ /$H^\gamma$ (y-axis) correlation. Each $H^N$ strip is assigned to a residue. The 2D NOESY correlates resonances that are close in space (< ~6 Å). The highlighted peak is a NOE between the $H^\gamma$ atoms of residue E445 of the tubulin main chain and the $H^N$ atom of the synthetically added $^{14}$N-Glu residue. (b) NOE highlighted in part (a) shown on the structure of the peptide.

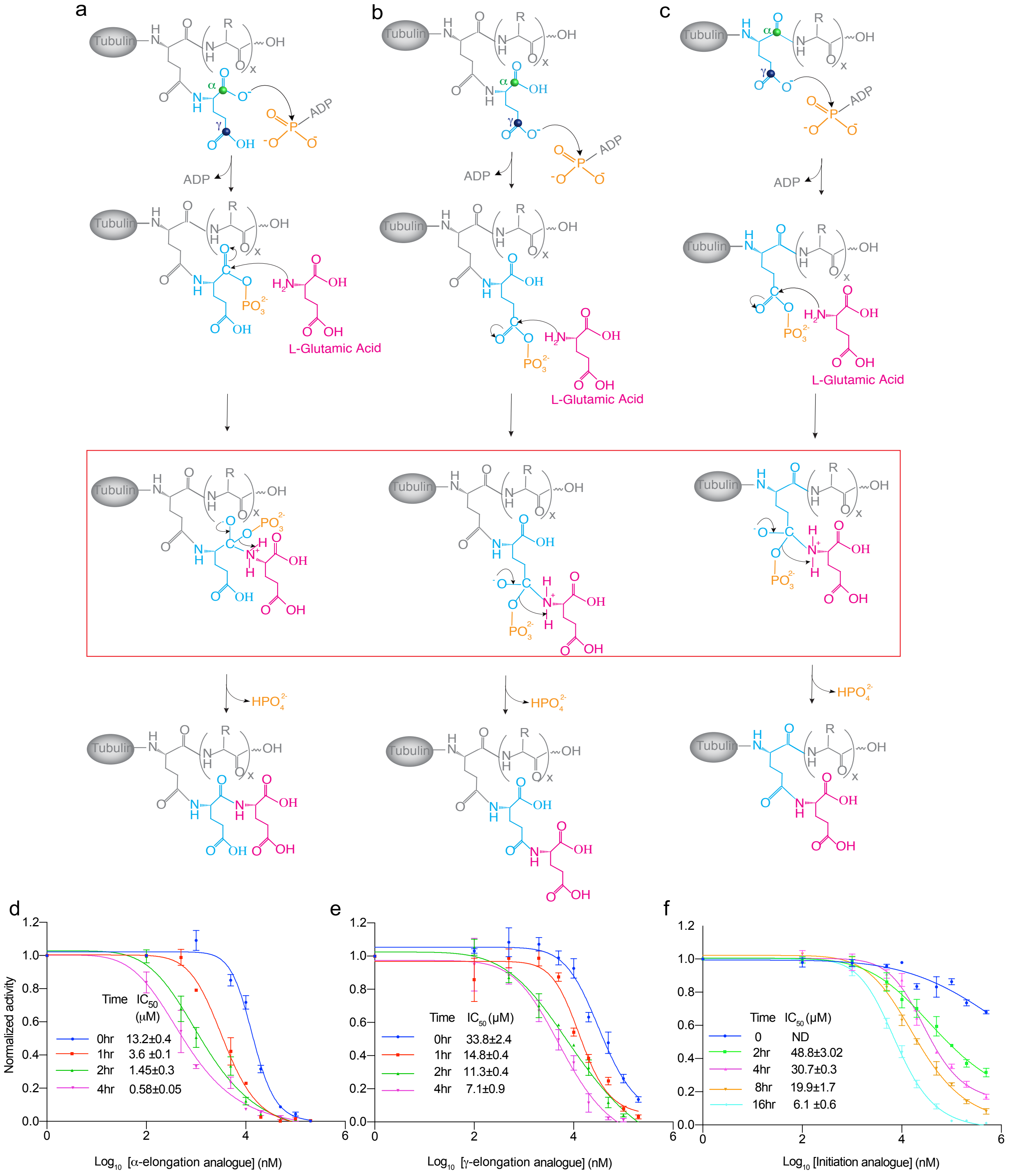


**Supplementary Figure 5. Proposed catalytic mechanism for TTLL6 and inhibition curves for the α-elongation, γ-elongation, and initiation analogs.**

(a-c) Proposed catalytic mechanism for(a) α-elongating, (b) γ-elongating and (c) initiating glutamylation reactions catalyzed by TTLL6. (d-f) Time dependent inhibition of TTLL6 by an α-elongation analog (d), γ-elongation analog (e) and initiation analog (f). Red box highlights tetrahedral intermediate state. Error bars indicate s.e.m (n=4).

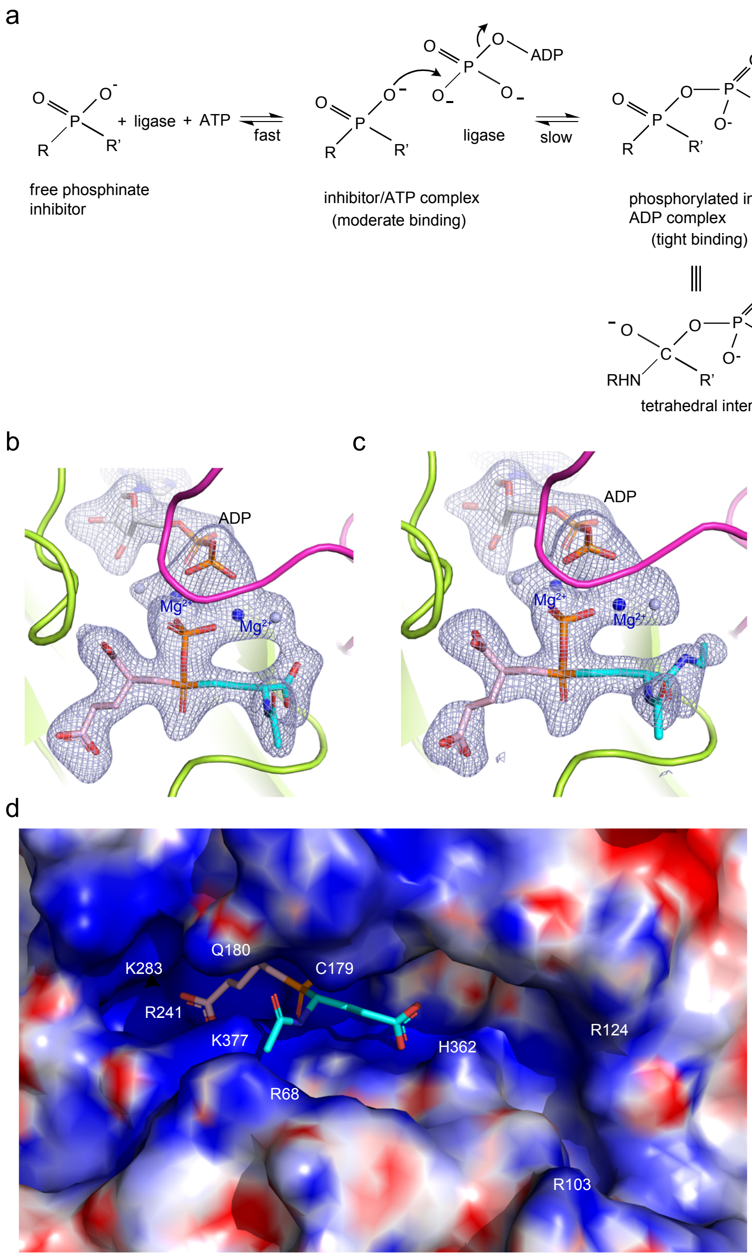


**Supplementary Figure 6. Structure of TTLL6 with tetrahedral intermediate analogs.**
(a) Mechanism for inhibitor phosphorylation (b) Active site showing the $|F_o|-|F_c|$ density (prior to modeling the $\alpha$-elongation analog) contoured at 4$\sigma$ (grey). (c) Active site showing the $|F_o|-|F_c|$ density (prior to modeling the initiation analog) contoured at 3$\sigma$ (grey).
(d) Electrostatic surface of the TTLL6 active site showing the electropositive character of the donor and acceptor glutamate binding site and the positively charged groove adjacent to the acceptor glutamate. $\alpha$-elongation analog shown as a stick model. The donor glutamate, transferred phosphate and acceptor glutamate of the $\alpha$-elongation analog are colored pink, orange and cyan, respectively. Conserved residues are labeled on the molecular surface.

a

| | | | | | | | | | | | | | |
|---|---|---|---|---|---|---|---|---|---|---|---|---|---|
| | | Glutamylase | | | | | | | | | Glycylase | | |
| | Initiase | ? | | | | | | | | | | | |
| | Elongase | ? | | | | | | | | | | | |
| TTLL6 | Function | TTLL1 | TTLL2 | TTLL4 | TTLL5 | TTLL7 | TTLL9 | TTLL11 | TTLL12 | TTLL13 | TTLL3 | TTLL8 | TTLL10 |
| R241 | H-bond with transferred phosphate | R | R | R | R | R | R | R | R | R | R | R | R |
| R219 | H-bond with transferred phosphate | R | R | R | R | R | R | R | R | R | R | R | R |
| K125 | H-bond with the β-phosphate of the ADP | K | K | K | K | K | K | K | K | K | K | K | K |
| D346 | $Mg^{2+}$ coordination | D | D | D | D | D | D | D | D | D | D | D | D |
| E359 | $Mg^{2+}$ coordination | E | E | E | E | E | E | E | E | E | E | E | E |
| K283 | Salt bridge with donor glu α-carboxylate | K | K | K | K | K | K | K | K | K | M | M | V |
| K377 | Salt bridge with donor Glu γ-carboxylate | K | K | K | K | K | K | K | - | K | C | C | I |
| C179 | Van der Waals interaction with carbonyl of acceptor Glu in γ-elongation analog | A | S | A | S | A | S | C | A | C | S | S | N |
| Q180 | H-bond with carbonyl of acceptor Glu in α-elongation analog | Q | R | R | R | M | Q | Q | R | Q | R | R | Q |
| H362 | Salt bridge with non-reactive carboxylate of acceptor Glu | A | Y | I | L | R | A | A | F | H | A | S | A |

b

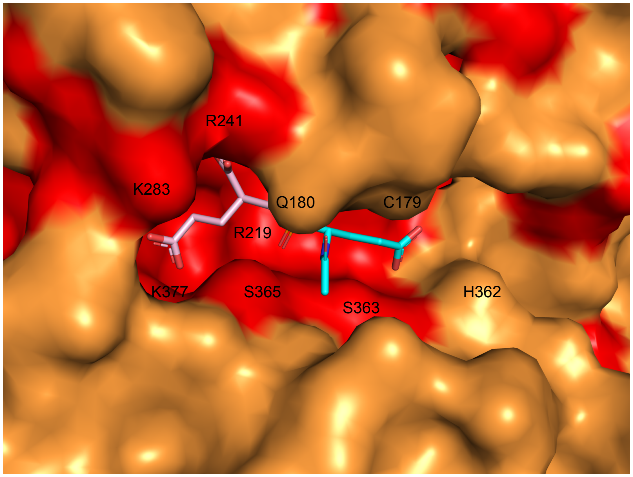


**Supplementary Figure 7 . Conservation of active site residues throughout the TTLL family.**
(a) Table showing conservation of critical active site residues among TTLL family members. Glutamylase initiases shown in grey, Glutamylase elongases in magenta; Glycylation initiases in green, Glycylation elongases in blue. (b) TTLL6 active site molecular surface color-coded according to conservation (as in (a)) illustrating the strong conservation in the donor glutamate binding site and variability of the acceptor glutamate binding site.

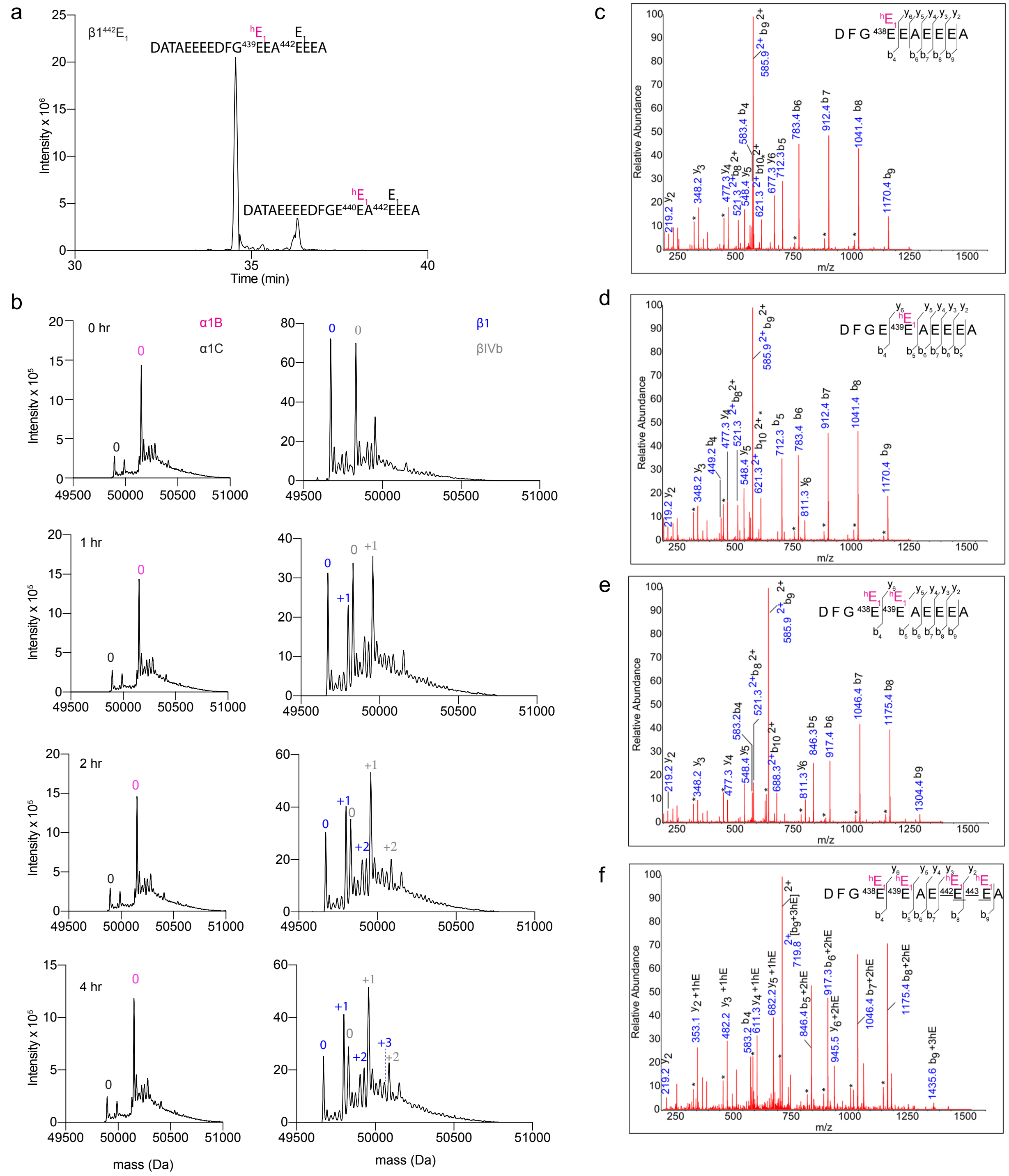


**Supplementary Figure 8. TTLL4 is a β-tubulin specific glutamylation initiase.**

(a) Extracted-ion chromatogram of enzymatically added monoglutamylated, $\beta 1^{442}E_1$ peptides with modifications at E439 and E440.

(b) Deconvoluted α- and β-tubulin mass spectra of Taxol stabilized human microtubules glutamylated by TTLL4 at a 1:10 enzyme:tubulin molar ratio after 0, 1, 2 and 4hr. The number of added glutamates is indicated and colored according to the tubulin isoform. (c-f) ) MS-MS sequencing of the β-tubulin C-terminal tails of microtubules glutamylated by TTLL4. Mono-glutamylated species are shown in (c) and (d), di-glutamylated species in (e), tri-glutamylated species in (f). Underline signifies that the spectra are ambiguous and the third glutamate can be added to either E442 or E443. Individual b, y-ion series and the amino acid sequence corresponding to each spectrum are indicated. Asterisks indicate ions with a neutral loss of a water molecule.

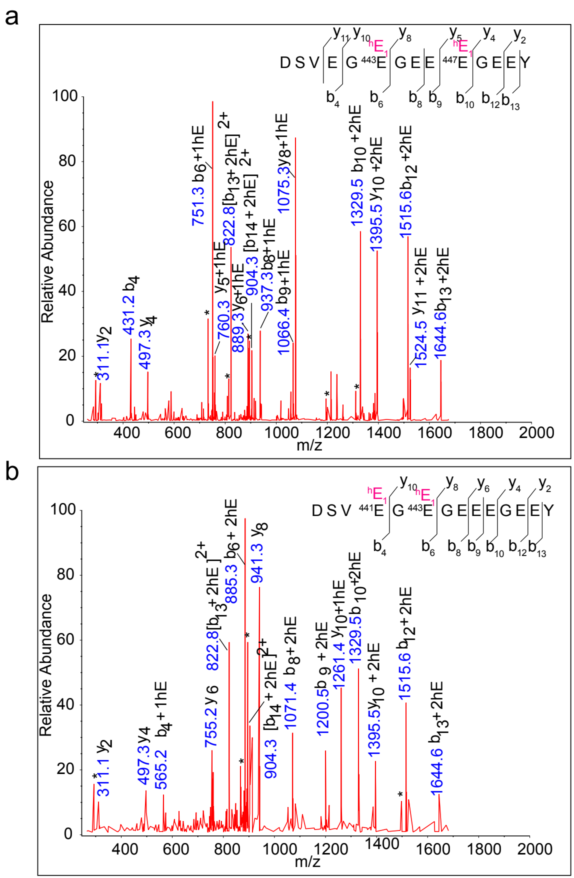


**Supplementary Figure 9. TTLL6 engineered mutant functions primarily as an initiase.**

MS-MS sequencing of the di-glutamylated $\alpha$-tubulin C-terminal tails of microtubules glutamylated by the TTLL6 C179A/Q180R/H362I mutant showing monoglutamylation at E443 and E447 (a) and E441 and E443 (b). Individual b-, y-ion series and the amino acid sequence corresponding to each spectrum are indicated. Asterisks indicate ions with a neutral loss of a water molecule.

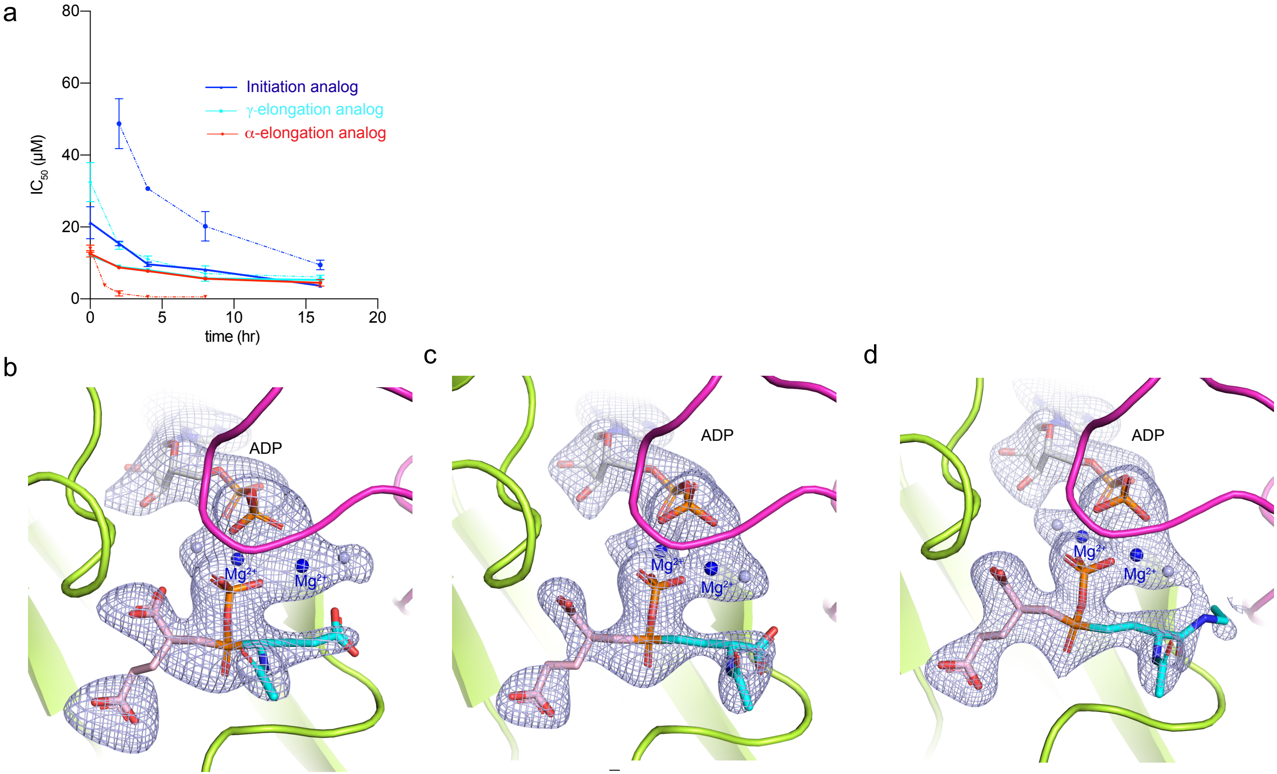


**Supplementary Figure 10. Inhibition curves and crystal structures for TTLL6 structure-based mutant with the α-elongation, γ-elongation, and initiation analogs.**

(a) Time dependent inhibition of TTLL6 C179A/Q180R/R182I/H362I/S367H mutant by the α-elongation, γ-elongation and initiation analogs. Error bars indicate s.e. of the fit (n=4). (b) Active site showing the $|F_o|-|F_c|$ density (prior to modeling the α-elongation analog) contoured at 3σ (grey). (c) Active site showing the $|F_o|-|F_c|$ density (prior to modeling the γ-elongation analog) contoured at 3σ (grey). (d) Active site showing the $|F_o|-|F_c|$ density (prior to modeling the initiation analog) contoured at 3σ (grey).
